# Flexible sensitivity to inputs during skilled tongue movements

**DOI:** 10.1101/2025.05.18.654702

**Authors:** Rajan Dasgupta, Mingyuan Dong, Daniel H. O’Connor

## Abstract

A hallmark of complex goal-directed movement sequences is the ability to rapidly switch motor programs by integrating incoming sensations with action context^1,2^, yet the underlying neural implementation remains elusive. Here, we demonstrate a network mechanism for flexibly adjusting the sensitivity of ongoing motor execution to external inputs in different sensorimotor contexts. We trained mice to make sequences of licks directed at a moving target. In a random subset of trials, they made “backtracking” licks by abruptly switching motor programs based on tactile feedback. We divided sessions into alternating cued blocks of trials; sensory-driven backtracking was required during one block-type, but not in the other. Targeted optogenetic stimulation of tongue/jaw somatosensory cortical inputs to the tongue premotor (anterolateral motor, or ALM) cortex that were precisely timed to arrive near the motor switching point reliably induced licking movements resembling those in backtracking trials. This effect was more readily induced during blocks with sensory-driven backtracking and was accompanied by larger optically-evoked deviations in neural activity. Population activity patterns could be separated along a latent axis that discriminated between block-types. On single trials, the location of population activity along this axis correlated with the impact of optogenetic stimulation and influenced the speed of sensory-driven motor switching. Our findings provide causal evidence for how external inputs are integrated with internal context signals to achieve flexible motor control.

---

We are uncannily proficient at carrying out sensory-guided motor tasks towards a goal. Driving behind a car, for example, requires one to constantly and precisely control the pressure on the throttle pedal, guided by visual input, with the goal of maintaining a safe distance. In addition, we must preserve the ability to make drastic changes (such as moving the foot to the brake pedal) given the appropriate salient stimulus (the brake-lights of the leading car turning on).

Finally, sudden braking would be inappropriate if the car is too far ahead. Thus, motor planning centers must simultaneously keep track of task context (“is the leading car too far?”), monitor the appearance of unpredictable stimuli (“did their brake-lights turn on?”) and maintain an internal estimate of continuous task variables (“how far is the car?”). How the brain solves such complex problems is a fascinating yet understudied subject.

Volitional movements have long been known to be closely intertwined with sensory feedback. Cortical motor areas integrate sensory inputs, context and goal information to produce dexterous movements^2–12^. This is true even for movements executed at relatively fast latencies, which can be influenced by intent and other contextual cues^13,14^. Fine motor control is critically reliant on crosstalk between a number of cortical and subcortical areas^8,15^. Although the importance of external inputs in motor planning is well established, the nature of its influence is underexplored and the underlying network mechanisms remain unclear. One issue complicating interpretation is that tuning properties of cortical single units can be notoriously complex^16^. More recently, alternate strategies that consider the activity of the recorded neural population as a whole have had much success. For instance, such approaches have revealed a rich repertoire of dynamics inside information-rich subspaces^17^ and provided neural computational underpinnings for a wide variety of observed behaviors^18,19^. Here, we coupled these analytical techniques with new experimental tools that enable dense electrophysiological recordings in the mouse orofacial sensorimotor system. This sophisticated yet tractable system has become a popular target to interrogate long-standing questions about how motor cortices carry out intricate sensory-guided movements^20^. In particular, ALM has emerged as a central player in planning and executing goal-directed orofacial movements^21,22^.

We trained mice to carry out a “sequence licking” task^23^ that relies on a cortical network including ALM and the tongue/jaw regions of the primary somatosensory cortex (tjS1) and primary motor cortex (tjM1)^24^. In each trial, mice made a series of directed licks towards a motorized lick port, thereby modulating their target tongue angle from left to right. The lick port unpredictably moved backwards (“backtracked”) in a random subset of trials, at a pre-trained branch-point, forcing the mice to monitor tactile input from the tongue and make rapid sensory-guided reconfigurations of ongoing motor execution. We divided sessions into alternating blocks of trials with either a 0 or 60% chance of backtracking, thus imposing a task context that signaled the need for motor planning flexibility. Surprisingly, we found that synaptically perturbing subpopulations of neurons in ALM at the appropriate point in the sequence, via optogenetic stimulation of inputs from tjS1, triggered backtracking-like behavior and neural population dynamics. Block context was encoded in all three cortical regions. In ALM, stronger block encoding predicted faster backtracking and greater susceptibility to optogenetic triggering of backtracking. Our findings reveal that the brain regulates the input sensitivity of motor cortex dynamics to achieve flexible, sensory-guided reconfiguration of behavior.

## A context-dependent sequence licking task with sensory-driven motor switching

We trained head-fixed mice to carry out sequences of directed licks in the dark following the delivery of an auditory go cue, towards a motorized lick port that moved through a series of target locations (Fig. 1a; Methods). Movements of the lick port to the next location in the sequence were triggered by tongue touches, so failures to touch (missed licks) resulted in the lick port remaining stationary. Once the final target location in the trial was licked, a water reward was delivered via the lick port. We monitored the mouse’s behavior at 400 Hz using high-speed videography and quantified post hoc the angle of the tongue with respect to the midline (“θ”, Fig. 1a) and its length (“L”).

**Fig. 1.**
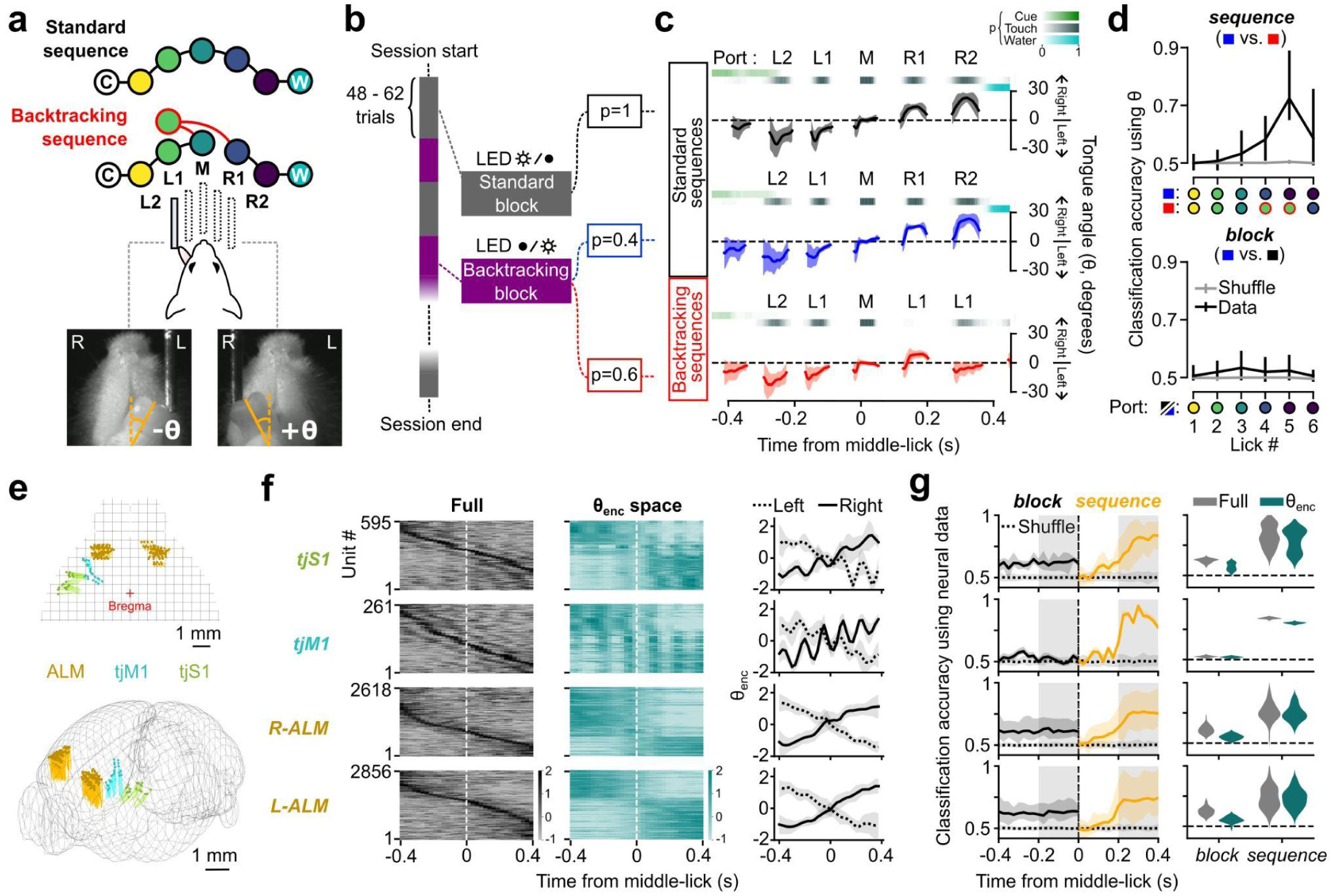
Context-dependent sequence licking task and neural encoding of task variables **a,** Top, schematic of sequences in the task. Following an auditory cue (“C”), the lick port was driven by touches from left to right. A touch at the final position triggered a water delivery (“W”). In backtracking sequences, the lick port moved backwards by one step following the middle-lick (“M”). Bottom, high speed video frame with tongue landmarks labeled by a CNN. Tongue angle (θ) was measured as deviation of the tongue midline from the vertical. The left (“L”) and right (“R”) sides of the mouse are indicated. **b,** Schematic of task structure. Blocks were cued to the mice with an LED. **c,** Mean θ from the three trial types in one example session, centered on time of middle-lick. Heatmaps denote probability of cue (green), touch (gray) and water (cyan). Error shading denotes s.e.m. **d,** Classification accuracy of trial type using θ for the first six licks. Sequence classification discriminated standard and backtracking sequences from Backtracking blocks, whereas block classification discriminated standard sequences from the two blocks. Colored circles indicate port location during each lick. Data are presented as mean ± 95% hierarchical bootstrap confidence intervals. **e,** Top, schematic of neural recordings. Bottom, approximate location of recording locations. **f,** Activity of single units in different regions during execution of standard sequences, centered on time of middle-lick. Left, heatmap of normalized firing rate PETHs of units sorted by time of peak of activity. Middle, heatmap of normalized activity in the θ_enc_ subspace (see Methods). The ordering of units is the same as the left heatmap. Right, mean θ_enc_ activity of units subdivided into two groups based on whether θ_enc_ activity was higher when the mouse licked to the left (i.e., before the middle-lick) or to the right (i.e., after the middle-lick). **g,** Classification accuracy of trial type using population neural activity recorded from the cortical regions in (f). Left, classification accuracy using full data versus time from middle-lick. Bins before the middle-lick were used for block classification; those after the middle-lick were used for sequence classification. Data are mean ± 95% hierarchical bootstrap confidence intervals. Gray shaded boxes indicate bins used for plots on the right. Right, violin plots of mean classification accuracy for block or sequence classification using full or θ_enc_ data.

In each trial, the lick port moved through one of two possible sequences. In the majority of trials in a session (70.8 ± 4% of trials, mean ± s.e.m., 69 sessions from 10 mice), the lick port progressed from left to right in an arc through a predictable sequence of five equidistant positions (“standard sequences”). In the remaining subset of trials, chosen at random, the lick port moved backwards by one location following touch at the middle position (“backtracking sequences”). This forced the mouse to make an abrupt adjustment of its ongoing motor program, based on sensory input from the tongue, to lick backwards towards the left in order to recover the sequence (Fig. 1a).

To investigate how task context affects motor planning, we designed the task to allow a variable prior expectation of encountering a backtracking sequence. Each session was composed of alternating blocks of trials, termed “Standard blocks’’ or “Backtracking blocks” (6.3 ± 0.1 blocks per session, 56.6 ± 0.2 trials per block, mean ± s.e.m., 69 sessions from 10 mice). Standard blocks contained exclusively standard sequences (Fig. 1b); thus a trained mouse could know with certainty that the lick port would move in a standard sequence. Backtracking blocks, on the other hand, contained a random mixture of standard (40% of trials) and backtracking (60%) sequences (Fig. 1b). Given the high likelihood of encountering backtracking sequences, we reasoned that the cortical θ-control programs in Backtracking blocks might be structured to facilitate faster motor switching based on sensory input. We explicitly signaled the current block-type to the mouse via an LED positioned in front of its face, which was either on for the entire duration of Standard blocks and off during Backtracking blocks or vice versa (n = 5 mice each, randomized across mice).

Mice learned to rapidly modulate the lick angle after training and were able to complete standard sequences within a second (0.99 ± 0.01 s per trial, median ± s.e.m., 15876 trials from 69 sessions and 10 mice; Fig, 1c). In backtracking sequences, mice typically encountered “missed” licks (in which the tongue failed to contact the lick port) by 0.2 s after the middle lick and learned to rapidly branch their motor programs in response, licking leftwards to contact the lick port and recover the sequence (Fig. 1c-d). Even though the majority of trials in Backtracking blocks required backtracking, mice did not backtrack by default, but instead tended to continue licking in a standard sequence pattern after the middle-lick until they encountered a missed lick at the following location (R1 in Fig. 1a) (Extended Data Fig. 1a). Thus, mice relied on tactile input at R1 to trigger backtracking.

We first asked whether the behavior of the mice reflected the current block-type and motor branching expectation. Since our task required mice to modulate target θ in a self-paced manner, block differences could either manifest as small dissimilarities in θ during individual licks, or in the gaps between successive licks. We reasoned that any potential differences were likely to show up after the middle-lick, around the time of motor switching uncertainty. Mice modulated θ in a highly stereotyped manner across licks in standard sequences, irrespective of which block they came from (Fig. 1c). Linear support-vector machine (SVM) classifiers failed to distinguish between individual standard sequence licks from different blocks above chance levels, even though similar classifiers could distinguish backtracking licks from standard licks (Fig. 1d). Mice did, however, slightly slow their lick rate after the middle-lick during Backtracking block standard sequences, when the mice were cued to expect potential backtracking (Extended Data Fig. 1b; inter-lick interval after middle-lick: 210 ± 2 ms, n = 19138 for Standard block standard sequences, 242 ± 7 ms, n = 7372 for Backtracking block standard sequences, mean ± s.e.m., 69 sessions and 10 mice)^25^.

## Block-type decoding from neural activity

We used four-shank Neuropixels 2.0 probes to record activity separately from multiple task-relevant cortical regions identified by stereotactic coordinates (Fig. 1e). In total, we recorded 595 single units from 5 sessions and 3 mice in left tjS1, 261 units from 5 sessions and 3 mice in left tjM1, 2618 units from 14 sessions and 8 mice in right ALM (R-ALM), and 2856 units from 14 sessions and 8 mice in left ALM (L-ALM). Activity of recorded units tiled the entire duration of sequence execution, with different units showing peak firing rates at different times in a trial. Units in ALM tended to show a slight preference for licking to the contralateral side (Fig. 1f).

Activity in large neural populations can be concentrated within a much smaller number of information-rich latent dimensions^1,26–28^. Moreover, a significant portion of the patterns of activity isolated via unsupervised dimensionality reduction methods may be ascribed to task-irrelevant movements and behaviors^29,30^. To interrogate task-associated dynamics within the population activity with greater power, we identified sets of latent orthogonal dimensions in the neural activity state space of each session that were best at decoding θ (Fig. 1f, Extended Data Fig. 2a-c; Methods). We refer to the subspace formed by these θ-correlated dimensions as the θ-encoding, or θ_enc_ space. In addition to considering the full dimensional data, we focused our analyses on activity projected onto θ_enc_, as it reflected patterns of activity that were widespread in the recorded population (Extended Data Fig. 2d, f) and helped highlight salient features of task-related neural dynamics. For instance, θ_enc_ space activity in different cortical regions showed qualitative differences in lick-to-lick fluctuations. Recapitulating our previous results^23^, θ-encoding in sensory and primary motor cortices showed rhythmic patterns correlated with individual lick cycles (Fig. 1f). These were notably absent in the θ_enc_ space within ALM activity, which tended to have smoother variation, consistent with continuously updating the target θ over the course of the sequence (Fig. 1f). θ_enc_ space activity also made prominent the licking side preference of individual units, with visible demarcations between left- and right-preferring units.

We asked whether the current trial-type could be decoded from the population neural activity during sequence execution (Fig. 1g and Extended Data Fig. 1c). For each time-bin, we trained Linear-SVMs to discriminate between either (i) Backtracking block standard sequence trials vs Backtracking block backtracking sequence trials (“sequence” classification), or (ii) Standard block standard sequence trials vs Backtracking block standard sequence trials (“block” classification). Sequence-type could be linearly decoded above chance levels from the population activity in all brain regions starting ∼0.2 s after the middle-lick (Fig. 1g). Sequence classification was most reliable in tjM1, consistent with a role as a driver of downstream motor output. We could also reliably decode the current block-type from activity sampled in most cortical regions. In contrast to sequence decoding, however, this was least accurate in tjM1 (Extended Data Fig. 1c). Block encoding strength, measured as the models’ block classification accuracy, was uniform throughout the trial, suggesting that task context is encoded in the population activity as a variable that stays constant during sequence execution. Interestingly, block-type could even be discriminated from the much lower dimensional θ_enc_ space (Fig. 1g).

This suggests that θ was encoded differently in the two block types and that the θ_enc_ space was also informative of other behavioral variables. Thus, although θ was indistinguishable in standard sequences from the two types of blocks, the underlying neural activity was not, suggesting that distinct motor programs mapped onto the same behavioral output.

## Activation of sensory cortical inputs to ALM evoked motor switching

Motor cortical dynamics during dexterous movements, like those of our licking task, are thought to be shaped by continuous sensory inputs to motor cortical areas^8,31^. To investigate the role of sensorimotor communication in our task, we adopted a strategy to optogenetically activate sensory cortical inputs to motor areas, particularly ALM. We sought to test two hypotheses: (i) ALM activity would be highly sensitive to long-range corticocortical inputs during sequence execution, and so optogenetic perturbation would produce large behavioral effects, especially in θ, and (ii) because our task required sensory-driven motor switching specifically in Backtracking blocks, ALM neural activity would be more perturbable in Backtracking block trials, such that the same optogenetic stimulus would lead to larger or more prolonged effects.

We targeted our optogenetic stimulation spatially to ALM-projecting sensory neurons using a combination of functional imaging and stereotactic-guided viral injections (Fig. 2a; Methods). First, we labeled motor cortex-projecting neurons by injecting (i) an adeno-associated virus (AAV) vector carrying Cre-dependent ChrimsonR-tdTomato in a wide area near tjS1, and (ii) a retroAAV construct carrying Cre-recombinase in ALM. Next, we localized tjS1 more precisely in each hemisphere by monitoring the intrinsic optical signals generated by the delivery of tactile stimuli to the tip of the tongue. Optical stimuli (595 nm, 40 Hz sinusoidal) were then delivered via optic fibers to the region where tdTomato expression and intrinsic signal responses overlapped. Although this procedure was used to label projection neurons bilaterally, optogenetic stimulation was applied to one side at a time.

**Fig. 2.**
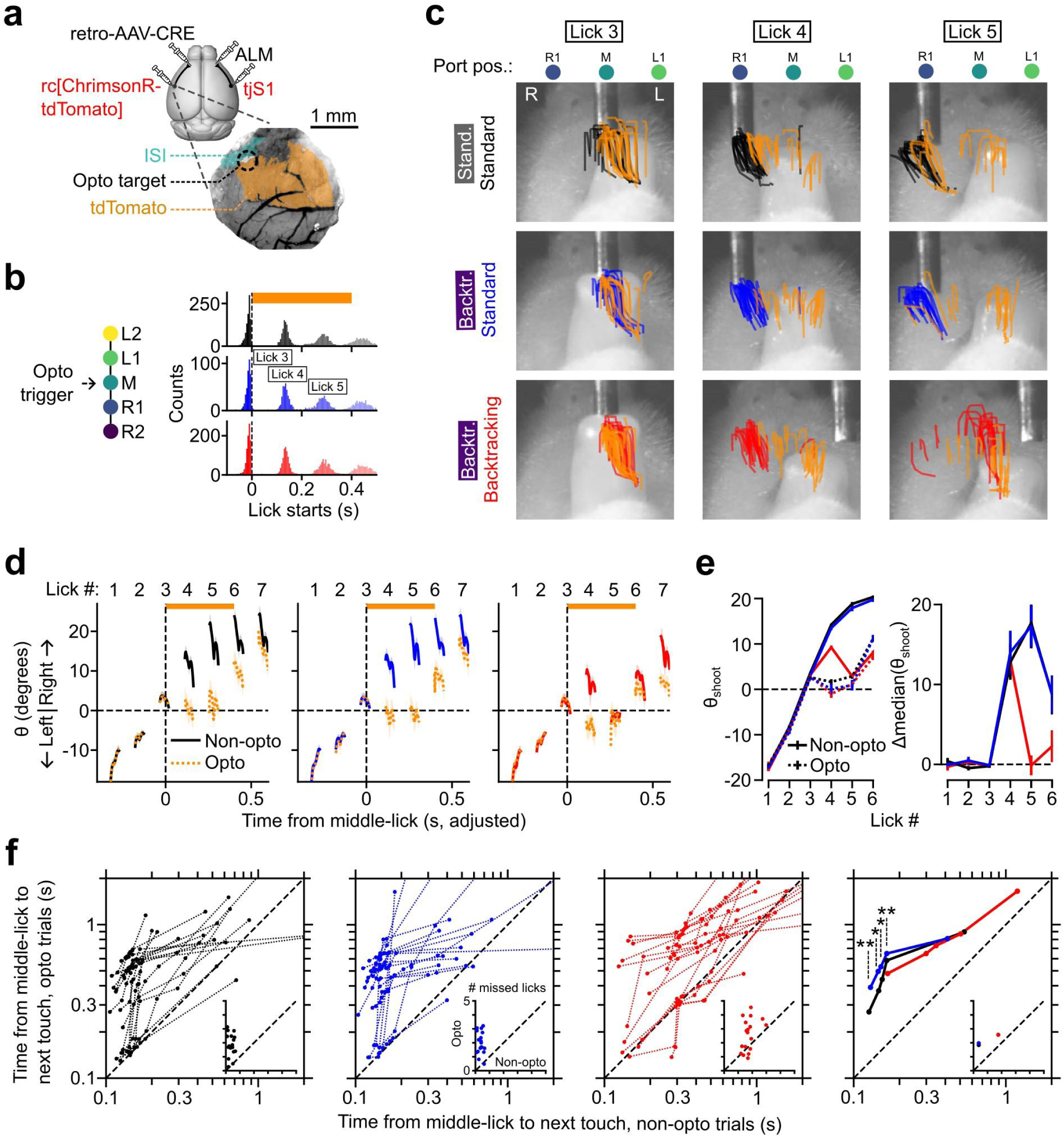
Optogenetic triggering of backtracking-like licks **a,** Top, schematic of viral injection strategy. Bottom, example image of cranial window over S1 showing locations of tdTomato labeling, ISI signal and target location for optogenetic stimulation. **b,** Left, schematic showing closed loop targeting of optogenetic perturbation to time of middle-lick. Right, histogram of lick initiation times relative to time of middle-lick in Standard block standard (black), Backtracking block standard (blue) and Backtracking block backtracking (red) trials. Orange bar indicates time of optogenetic stimulation. Lick numbers in boxes indicate which lick in the sequence each peak corresponds to. **c,** Tongue tip traces of single licks from trials with (“opto”) and without (“non-opto”) optogenetic stimulation in an example session. Black, blue and red traces show non-opto trials corresponding to the same trial types as (b). Overlapping orange traces show licks from opto trials of the corresponding type. The left, middle and right columns show tongue tip traces for licks 3, 4 and 5 respectively. The background images in each row are high speed video frames from example opto trials taken during the execution of corresponding licks. The left (“L”) and right (“R”) sides of the mouse’s face and the relevant port positions (pos.) are indicated. **d,** θ versus time from middle-lick for the three trial types. Orange bar denotes time of opto-stimulation. Data are mean ± s.e.m. **e,** Left, θ_shoot_ (see Methods) versus lick number for non-opto and opto trials. Right, difference in median θ_shoot_ values between non-opto and opto trials, for each trial-type and each session. Positive values indicate opto-induced deviations to the left. Data are mean ± s.e.m. **f,** Quantile-quantile plots showing time between middle-lick and next touch for opto versus non-opto trials. The left three plots show values for each individual session for each of the three trial types (same colors as in (b)), with data points from the same session connected by dotted lines. The right plot shows medians across sessions. Shown are the 0th, 25th, 50th, 75th and 100th percentile values for each session. Insets are scatter-plots showing the mean number of missed licks between middle-lick and next touch for opto versus non-opto trials. One-tailed paired t-test with the alternative hypothesis being that blue y-axis quantiles are greater than black y-axis quantiles, ** p < 0.01, * p < 0.05.

To examine the sensitivity of ALM population activity to external inputs, we timed the optogenetic stimulus to coincide with the point in the sequence when sensory inputs were most behaviorally relevant, i.e., immediately preceding the potential motor branchpoint. The LED stimulus was triggered in a closed loop manner by the tongue touching the lick port at the middle location (Fig. 2b, left), and lasted 400 ms. Given the roughly 160-ms lick cycle period, this stimulation paradigm allowed us to target the two licks that immediately followed the middle-lick (Fig. 2b, right), which were lick numbers 4 and 5 in the full sequence. In backtracking sequences, these licks typically corresponded to a missed lick (targeted towards the R1 location) followed by a backtracking lick (targeted towards the port at L1, Fig. 1c, bottom row, red traces).

We triggered the optogenetic stimulus LED in a random subset (25-30%) of trials (“opto” trials) while mice executed sequences. Intriguingly, unilateral optogenetic activation of tjS1→ALM neurons evoked licks that resembled backtracking (Fig. 2c-e). In all mice, both of the licks targeted by optogenetic stimulation (lick numbers 4 and 5; orange traces in Fig. 2c) tended to be directed towards the left compared to the corresponding non-opto licks. The magnitude of the opto-induced deviation in θ varied from trial to trial but was readily apparent when aggregated across trials (Fig. 2d). Unilateral inactivation of ALM can produce prominent ipsilateral θ biasing effects^32^. In our task, unilateral ALM activation on either side produced licking towards the left (Extended Data Fig. 4a-b). For standard sequences from either block-type, optogenetic perturbation produced large changes in the tongue protrusion angle (θ_shoot_) in both of the targeted licks (Fig. 2e). For backtracking sequences, optogenetic stimulation produced similarly large deflections in θ_shoot_ (Δθ_shoot_) for the first targeted lick (lick 4). But because lick 5 in non-opto backtracking sequences was also typically directed towards the L1 location, tongue traces in trials with and without optogenetic stimulation overlapped during this lick (Fig. 2c-e), leading to a reduced Δθ_shoot_ during the second half of the opto-stimulus period (Fig. 2e).

We wondered whether the stimulation of somatosensory cortical inputs to ALM from regions outside the tongue/jaw area could be sufficient to induce backtracking-like licks. In separate sessions, we delivered optical stimulation to cortical regions that did not show an ISI response (ISI**^-^**), but nevertheless had tdTomato expression, indicating the presence of ALM-projecting neurons (Extended Data Fig. 3a-b). ISI**^-^** stimulation at middle-lick also produced backtracking-like licks (Extended Data Fig. 3c-d). Although some indirect activation of tjS1 neurons could not be ruled out, this suggested that stimulation need not be restricted to tjS1→ALM inputs in order to evoke backtracking-like licks. However, ISI**^-^** stimulation was less robust than ISI^+^ stimulation at inducing backtracking, as evidenced by a reduced effect at lick 5 and faster recovery following offset of the opto stimulus (Extended Data Fig. 3d), suggesting that the tjS1→ALM pathway was uniquely potent at producing the effect.

Licks produced during LED stimulation also tended to be hypometric (Extended Data Fig. 4c). Opto-evoked backtracking-like licks, while similar in tongue protrusion angle to normal backtracking licks, thus failed to reach the lick port in most trials, hindering behavioral performance even during backtracking sequences (Fig. 2d, f). The probability of licking was also reduced during optogenetic stimulation (Extended Data Fig. 4d; Methods).

Mice quickly recovered their behavior after the end of the optogenetic stimulation period, directing their next lick towards the location where they would have licked in the absence of the LED stimulus (Fig. 2d). If the expectation of backtracking in Backtracking blocks made ALM activity easier to disrupt, then we should expect the mice to take longer to recover their behavior following optogenetic stimulation during those blocks. To test this, we quantified (i) the number of erroneous licks and (ii) the gap in time between the middle-lick (i.e., the start of the LED stimulus) and the next lick port contact. Optogenetic stimulation increased both the number of missed licks and the time between successive lick port contacts in all trial types and in most sessions (Fig. 2f). This effect was slightly larger for standard sequences occurring in

Backtracking blocks compared with Standard blocks (Fig. 2f, right), indicating that sequence progression was indeed more perturbable when mice expected backtracking sequences. This block-type difference could not be explained by the slowing of lick cycles that otherwise occurs in non-opto trials after the middle-lick (Extended Data Fig. 4g). ISI**^-^** optogenetic stimulation increased the number of erroneous licks and the time gap to a lesser extent (Extended Data Fig. 3e).

In control experiments we triggered the optogenetic stimulation (10% of trials) at the time of water delivery, when tactile input was no longer needed to decide whether to backtrack. This failed to evoke backtracking-like licks and did not cause significant changes in the lick rate during the period of stimulation (Extended Data Fig. 4e-f). Thus, sequence behavior was specifically perturbable during the period when touch was most relevant.

Together, these experiments suggest that appropriately timed sensory cortical input to ALM could trigger switching from an ongoing to an alternative motor program.

## Optogenetic perturbation drove population activity to motor-switching states

We next investigated the effects of optogenetic perturbation on ALM neural population activity. We recorded from ipsilateral ALM during optogenetic activation of tjS1→ALM neurons (n = 14 sessions from 4 mice). Optogenetic stimulation evoked short latency spikes in a small fraction of ALM units (Fig. 3a; out of 1743 units, 122 units increased their mean firing rates by at least 1 s.d., 63 by at least 2 s.d. and 40 by 3 s.d. in the first 25 ms of optogenetic stimulation, relative to non-opto trials), indicative of monosynaptic connections between tjS1 and ALM neurons. A number of units were also inhibited (out of 1743 units, 31 units decreased their mean firing rates by at least 1 s.d. and 6 by 2 s.d. in the 50-100 ms period from the start of optogenetic stimulation, relative to non-opto trials). Optogenetic stimulation evoked dramatic changes in neural population trajectories in the θ_enc_ space for all three trial-types, halting or pushing the trajectory backwards (Fig. 3b, leftward on the x-axes) as in normal, sensory-guided backtracking (Fig. 3b, right).

**Fig. 3.**
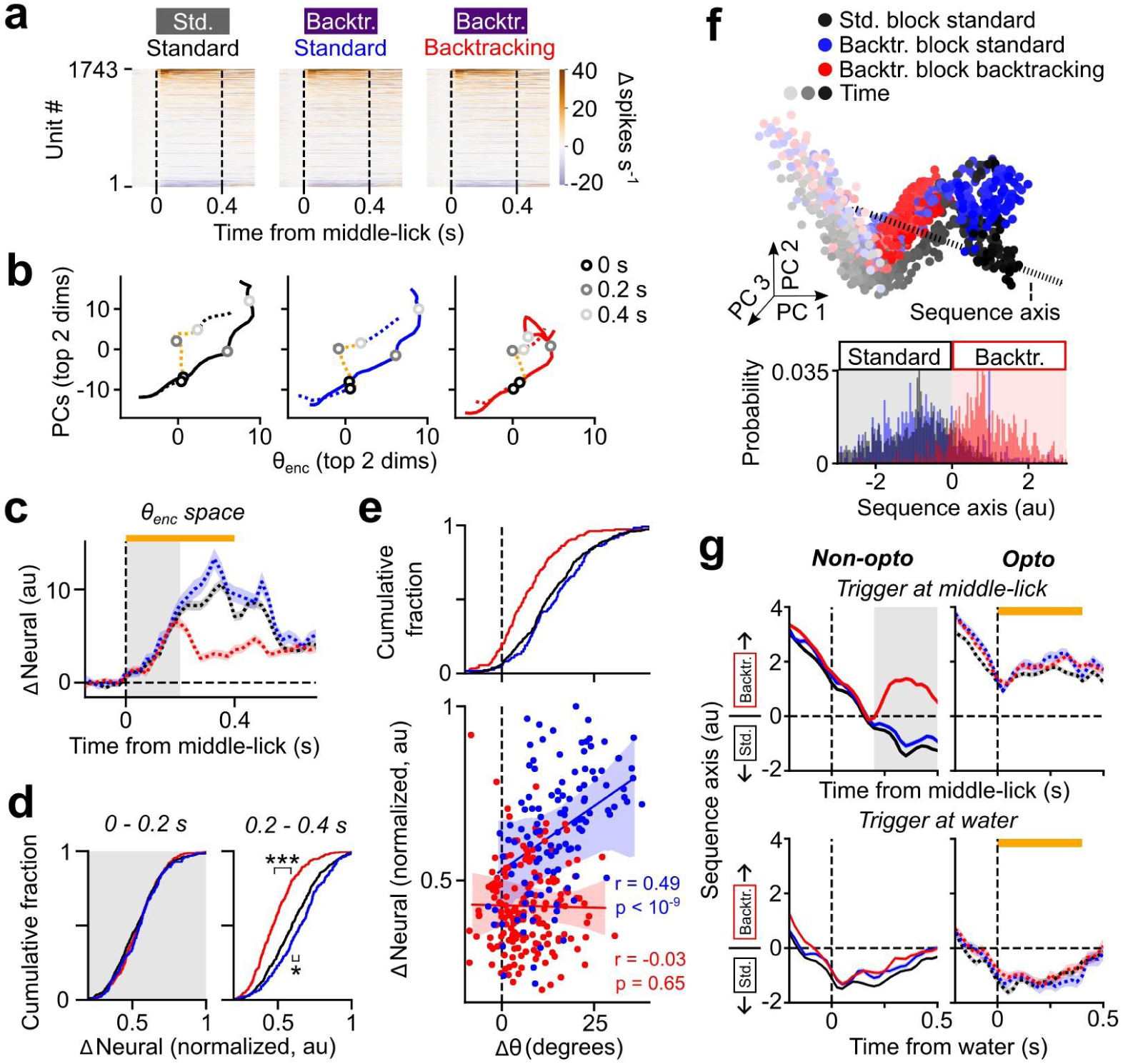
Optogenetic triggering of backtracking-like population activity **a,** Heatmaps showing change in firing rates for each ALM unit due to optogenetic stimulation in each of the three trial types. Dashed lines indicate start and end times of optogenetic stimulation. **b,** Mean activity in the top two principal components (“PCs”) versus the top two θ_enc_ dimensions for Standard block standard (black, left), Backtracking block standard (blue, middle) and Backtracking block backtracking (red, right) trials. Solid lines show mean of non-opto trials, dotted lines show mean of opto trials. Period of opto-stimulation is shown in orange, along with markers for 0, 0.2 and 0.4 s after middle-lick. **c,** Baseline-subtracted change in neural activity in the θ_enc_ subspace, versus time from middle-lick. Line colors correspond to the same trial types as in (b). Each opto trial was compared to the median of the corresponding non-opto trials in the same session. Data are mean ± s.e.m. Orange bars denote time of opto-stimulation. Gray shading area delineates halves summarized in (d). **d,** Cumulative histograms showing mean normalized ΔNeural in the first half (0 - 0.2 s, left column) and second half (0.2 - 0.4 s, right column) of opto-stimulation for the θ_enc_ subspace. Two-sided Kolmogorov-Smirnov test, *** p < 0.001, * p < 0.05. **e,** Top, cumulative histogram of deviations produced in θ for the three trial types (same color scheme as in (b), Methods). Bottom, scatter-plot showing normalized ΔNeural versus Δθ for Backtracking block standard (blue) and backtracking (red) trials. Solid lines show mean linear fits to data, error shading represents 95% hierarchical bootstrap confidence intervals. **f,** Top, trials from an example session plotted in top 3 PCs. The identified sequence axis is shown. Bottom, histogram of trials projected onto the sequence axis. More positive values indicated activity consistent with backtracking sequences. **g,** Projections of trials onto the sequence axis versus time from middle-lick (top row) or water delivery (bottom row) for non-opto (left column) or opto (right column) trials. Data are mean ± s.e.m. Orange bars denote times of opto-stimulation. Gray shading in the top left plot indicates time bins used for sequence axis calculation.

We quantified the magnitude of the opto-induced perturbation in θ_enc_ space population activity as the bin-by-bin differences in the state space trajectories of each opto trial from the corresponding median non-opto trial (“ΔNeural”; Methods). These opto-induced deviations diverged for the different trial-types after ∼0.2 s (Fig. 3c). We therefore considered the two halves of the stimulation period separately. The magnitude of perturbation in the θ_enc_ space was similar for all three trial-types in the first half of the stimulation period (0 - 0.2 s from middle-lick; gray shading in Fig. 3c), but diverged in the second half (0.2 - 0.4 s from middle-lick, Fig. 3c-d). For backtracking sequences, ΔNeural decreased after ∼0.2 s (Fig. 3c, red trace). During execution of non-opto backtracking sequences, neural activity distinguishes backtracking at approximately 0.2 s after the middle-lick (Fig. 1g, left), so this decrease in ΔNeural for backtracking sequence trials is consistent with the hypothesis that optogenetic perturbation caused activity in the θ_enc_ space to resemble that seen during backtracking. In contrast, the ΔNeural for standard sequence trials increased in the second half of the optogenetic stimulation period (Fig. 3c). Optogenetic stimulation produced slightly larger values of ΔNeural during Backtracking block standard sequences compared with Standard block standard sequences (Fig. 3d, right), indicating that ALM is more susceptible to perturbation during Backtracking blocks.

If opto-induced deviations in θ were caused by opto-induced deviations in ALM activity, then trial-to-trial variability in the two should be correlated. Indeed, we found that the magnitudes of opto-induced perturbations in ALM neural activity and behavior were correlated for standard sequences (Fig. 3e, bottom).

To determine if optogenetic stimulation pushed the trajectories in a behaviorally meaningful direction in neural state space, we identified latent dimensions that were best at decoding the sequence type (Backtracking block standard vs. Backtracking block backtracking) in the period 0.2 s after middle-lick (Fig. 3f top; Methods). Projecting data onto this “sequence axis” led to separation of backtracking sequences from standard sequences of either block-type (Fig. 3f, bottom). Optogenetic perturbation at the middle-lick pushed activity towards the backtracking (positive) direction on the sequence axis in all trial-types (Fig. 3g, top), consistent with its behavioral effects (Fig. 2). This effect was absent when the same stimulation was delivered at time of water reward (Fig. 3g, bottom), despite a similar degree of modulation of individual single-units (Extended Data Fig. 4h). This suggested that licking motor output became uncoupled from perturbations of ALM dynamics upon sequence completion.

Together, these findings suggest that inputs to ALM could push its dynamics from those for controlling standard sequences to those for backtracking sequences, specifically at times near the sequence branch point.

## Optogenetic stimulation preferentially activated motor switch-preferring units

We next turned to the encoding of trial-types at the level of single units. We used ideal observer analysis to identify units that showed a significant preference in their firing rates prior to the middle-lick for either block-type, or after the middle-lick for either sequence-type (Methods).

Considering that backtracking represented a drastic reversal of the direction of θ-progression mid-sequence, our a priori expectation was to find a preponderant population of units that preferentially responded to backtracking-related sensory cues. Surprisingly, standard sequence preferring units greatly outnumbered backtracking sequence preferring units in all regions (Extended Data Fig. 5a), while units preferring either block-type were roughly equal in numbers. We reasoned that if the smaller proportion of backtracking-preferring units in ALM were still responsible for driving the switch to backtracking modes, they should show elevated firing in contexts where the likelihood of encountering backtracking trials was greater. Indeed, backtracking sequence preferring units tended to also prefer Backtracking blocks (and vice versa, Extended Data Fig. 5b, left) while Backtracking block preferring units showed a greater preference for backtracking sequences than those that preferred Standard blocks (Extended Data Fig. 5b, right). Moreover, units that showed higher θ_enc_ space activity when licking to the left (i.e., prior to the middle-lick in Fig. 1f, middle) tended to show a preference both for backtracking sequences and Backtracking blocks (Extended Data Fig. 5c). Thus, backtracking sequence preferring units comprised a relatively small sub-population within ALM that were primed to fire in Backtracking blocks prior to the middle-lick.

Optogenetic stimulation caused population activity in ALM to resemble backtracking-like states (Fig. 3). Consistent with this, we found that units that preferred backtracking sequences were more strongly activated and less inhibited by the optogenetic stimulation than units that preferred standard sequences (Extended Data Fig. 6a). Motivated by the observation that ALM activity during non-opto standard and backtracking sequences started to diverge around 0.2 s after the middle lick (Fig. 1g), we wondered whether the sub-population of units that drove this divergence overlapped with the population of units targeted by tjS1 inputs. We divided the population of units that showed a preference for either sequence-type into groups based on the time at which they started to show their sequence preference, and considered separately the impact of opto-stimulation on the activity of each group. Interestingly, units that started discriminating between sequence-types at the 0.2 s bin were more excited by the optogenetic stimulation than units that started showing sequence preference at other times (Extended Data Fig. 6b), suggesting that these units were especially sensitive to sensory inputs. Together, these findings suggested that the small subpopulation of backtracking-preferring units that were more active during Backtracking blocks were also preferentially excited by external inputs.

## Block-type encoding influenced motor switching performance

We next asked whether block-type encoding at the population level in sensorimotor cortical regions carried any significance for performing the task. Some of our findings suggested small but consistent increases in the perturbability of ALM activity in Backtracking blocks. If the cued expectation of sequence branching indeed had an impact on how easily backtracking could be induced, this should be reflected in the trial-to-trial variability of backtracking performance. To test this prediction, we identified the latent dimension that best separated the activity state spaces corresponding to the two block-types immediately prior to the sequence branch point (Fig. 4a). We termed this latent dimension the “block axis”. Projections of individual trials from Standard and Backtracking blocks onto this axis were separated about 0, with most Backtracking block trials being assigned negative values (Fig. 4b). We took the absolute value of the block axis projection on each trial (which gives the confidence that it came from one or the other block-type) to represent the awareness of the mouse about the current block (cf. ^18^).

**Fig. 4.**
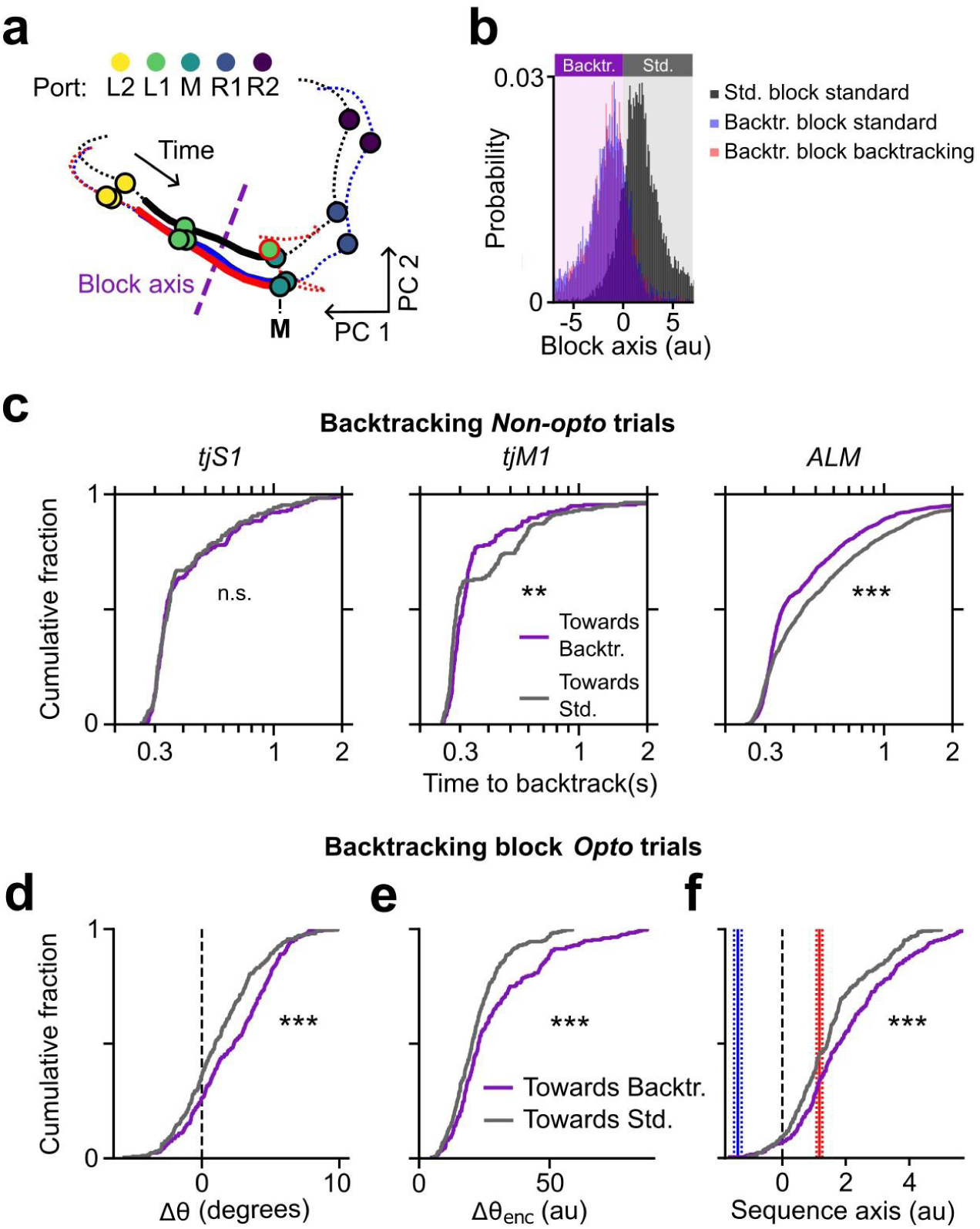
Block-encoding strength influenced motor switching performance and response to perturbation a, Mean activity in the top two PCs for Standard block standard (black), Backtracking block standard (blue) and Backtracking block backtracking (red) trials. Colored circles indicate median times of lick port contacts at different locations. Solid lines indicate time bins (250 ms) used to calculate block axis, plotted here for this session as a purple dashed line. **b,** Histogram of projections onto block axis for single trials (same color scheme as in (a). More negative values are indicative of activity consistent with Backtracking block trials. **c,** Cumulative histograms for each cortical region of time taken to backtrack (time between middle-lick and next touch) in non-opto backtracking trials. The data points were split by their median block axis projection value into more negative (purple, “Towards Backtr.”) and less negative (gray, “Towards Std.”) halves. **d,** Cumulative histograms of opto-induced deviations in θ for all opto trials in Backtracking blocks. Trials were split (similarly to c) by the median block axis values into those with more negative (purple, “Towards Backtr.”) or less negative (gray, “Towards Std.”) block axis projections. **e,** Cumulative histograms of opto-induced deviations in θ_enc_ activity for all opto trials in Backtracking blocks. Trials were split (similarly to c-d) by the median block axis values. **f,** Cumulative histograms of sequence axis projections of all opto trials in Backtracking blocks, split (similarly to c-e) by the median block axis values. Blue and red horizontal lines show mean sequence axis projections for non-opto Backtracking block standard and backtracking trials respectively (mean ± s.e.m.). Two-sided Kolmogorov-Smirnov test, *** p < 0.001, ** p < 0.01.

We reasoned that if stronger block-encoding represented greater prior expectation of encountering backtracking trials, we should observe faster sensory-driven motor switches to backtracking in those trials. To test this, we subdivided all non-opto backtracking trials by their median block axis projection value into two groups with stronger or weaker block encoding (i.e., trials whose projections were more towards or less towards the Backtracking block space on the block axis). Consistent with our hypothesis, we found that in trials that had more negative block axis projections prior to the middle-lick (“Towards Backtracking” trials, Fig. 4c), mice were able to more rapidly switch to backtracking in response to sensory input compared to trials that had more positive block axis projections (“Towards Standard” trials, Fig. 4c). Furthermore, trials in which mice successfully backtracked after a single missed lick showed stronger block-type encoding than trials in which they made multiple missed licks before backtracking (Extended Data Fig. 7). Interestingly, this trend was evident in data from motor (ALM, tjM1) but not sensory (tjS1) cortices (Fig. 4c), suggesting that information about the current block facilitated motor switching only in regions that directly drive motor output.

Next, we examined whether block-encoding strength influenced ALM perturbability by optogenetic stimulation of long-range cortical inputs to ALM. We subdivided Backtracking block opto trials by their median block axis projections and considered the magnitude of the opto-induced deviations in θ and θ_enc_ for each group. Similar to sensory-driven backtracking, trials closer to the Backtracking block space were perturbed to a larger extent by the optogenetic stimulation than trials that were closer to Standard block space (Fig. 4d, e). As optogenetic stimulation caused neural activity to resemble backtracking-like activity, we wondered whether block-encoding strength also correlated with this effect. Indeed, trials with stronger block encoding were also pushed more towards the backtracking direction (i.e., more positive values) on the sequence axis by the optogenetic stimulation (Fig. 4f). Thus, greater expectation of backtracking prior to the middle-lick enabled easier switches to backtracking-like modes after the middle-lick.

Finally, we examined whether population dynamics in the two block-types showed any qualitative differences. First, to characterize the population activity in each block, we identified a number of largely uncorrelated parameters of neural activity and quantified them separately for each trial (Extended Data Fig. 8a; Methods). Then we trained Random Forest classifiers to discriminate the block-type of each trial using these parameters and the block axis projections as features. We shuffled each individual parameter to gauge its importance to the classifiers’ performance. Block axis projection values were more crucial than any of the other parameters for block classification accuracy and were the primary criterion used by most individual trees in the ensemble to determine block-type (Extended Data Fig. 8b-c). This indicated that the nature of block encoding was a simple linear shift within the neural state space, rather than dramatically altered population trajectories.

To summarize, we trained mice to make rapid sensory-guided motor branching, or backtracking, during skilled motor execution and maintain an internal expectation of the likelihood of making such adjustments in different blocks. Spatiotemporally precise optogenetic perturbations of motor planning activity evoked backtracking-like behavior and activity, and expectation of backtracking led to larger optogenetic effects. Strength of block encoding influenced the speed of backtracking behavior. Together, these findings suggest that the sensitivity of motor cortical activity to external inputs is flexibly regulated (Extended Data Fig. 8d).

## Discussion

We found that activation of long-range cortical inputs to motor planning centers was unexpectedly potent at inducing sequence branching-like behavior and neural states (Figs. 3, 4, Extended Data Figs. 3, 4). This suggests that neural activity in ALM was primed to switch to backtracking modes, such that even relatively small synaptic perturbations of ALM populations were sufficient to push the system into backtracking-like cortical states, particularly around the time of the middle-lick when sensory input was most pertinent^33,34^. Indeed, the same optogenetic intervention had little impact on behavior when it was delivered at the end of sequence execution (Extended Data Fig. 4e, f), suggesting that external perturbations to the ALM somehow ceased to affect motor output when licking was no longer task-directed. Given that ours was a self-paced task, and assuming that it was the animals’ objective to maximize accumulated reward as a function of time, it makes normative sense to switch motor programs as soon as the sensory hallmarks of backtracking lick port movements are detected. However, accurate control of the backtracking trigger is a non-trivial problem for the brain to solve, as balance must be maintained between rapidly switching to backtracking and avoiding triggering it when uncalled for. One way to mitigate the problem could be to maintain an internal estimate of the likelihood of encountering a backtracking sequence and to adjust accordingly the potency of the priming of the backtracking trigger. We encouraged such an estimate by explicitly cueing the mice to expect backtracking in Backtracking blocks but not in Standard blocks. We tested an intuitive hypothesis: that execution of movements with higher chances of encountering backtracking would be more responsive to external synaptic inputs. We found that block information was widely encoded across cortical regions, and in the motor cortices, it influenced the speed with which the system could switch to backtracking. Thus, the sensitivity of ongoing cortical motor programs to external inputs was context-dependent, whether those inputs came in the form of behavioral cues or optogenetic perturbations.

Interestingly, θ during standard sequence execution was largely indistinguishable between blocks and major features of population activity were not different. Block-type encoding may thus be thought of as a conditional output-null factor^35,36^ - information that exists within the cortical activity, yet produces little readily observable impact on the motor output outside of the specific conditions in which that information becomes relevant, in this case inputs occurring near the motor branching point. A caveat to this interpretation is that we cannot rule out that the behavior of the mice differed between blocks in unobserved ways.

Our results offer insights into the larger theme of hierarchical models for the control of complex motor behavior. Generally, these models afford significant autonomy to lower level structures, such as the midbrain, striatum or superior colliculus, to control both the kinematics of individual movements and the selection of movement sequences^32,37–39^. In our task, mice were highly practiced at performing lick sequences. This was true also for backtracking sequences, as mice encountered hundreds of backtracking trials by the time they were deemed ready for recording sessions. This repeated exposure to backtracking-related sensory cues allowed mice to become highly skilled at the ensuant motor response, which may have been learned and then cached^38^ to be engaged as needed, representing a form of amortized control^40^. The motor cortex, which we have previously shown is critical for our task^23^, mediates movement selection when it is guided by unpredictable stimuli^39,41,42^, such as those encountered in our backtracking trials.

Moreover, our task involved a contextual factor—block-type—that cued the need to respond to unpredictable stimuli. This represents an additional layer of complexity required of the sensorimotor control strategy necessary in our task. Our results show that flexible regulation of the input sensitivity of ALM is a key mechanism for achieving this context-dependent sensorimotor control.

## Acknowledgments

We thank Timothy Harris for equipment; Duo Xu, Varun Chokshi, William Olson and Yi-Ting Chang for suggestions and assistance; Michael Economo for discussion. This work was supported by NIH grants R01NS089652, 1R01NS104834-01, 1RF1NS131984-01, 1RF1NS131984-01, 1U19NS137920-01 and 1U01NS115587-01.

## Author contributions

R.D. and D.H.O. devised the study. R.D. and M.D. developed experimental protocols. R.D. conducted experiments, developed code and analyzed data. R.D. and D.H.O. wrote the manuscript.

## Methods

### Mice

All procedures were carried out in accordance with protocols approved by the Johns Hopkins University Animal Care and Use Committee (protocols: MO21M195, MO24M185). We obtained behavioral/neural data from 11 wild-type mice (C57BL/6J, JAX #000664, 7 female, 4 male) at 3-10 months of age. Mice were housed in a reverse light cycle room.

### Surgeries and intrinsic signal imaging

#### Headpost implantation

For implantation of titanium headposts, mice were anesthetized with isoflurane (1-2% in O_2_; Surgivet). The scalp was excised, the periosteum was removed, and the skull was scored. A layer of cyanoacrylate adhesive (Krazy Glue) was applied. The headpost was affixed on top using either dental acrylic (Lang Dental) or Metabond.

#### Viral injections and cranial windows

In a subset of mice, we virally expressed a light-gated opsin in ALM-projecting tjS1 neurons. First, we made bilateral craniotomies (2.5 - 3 mm diameter) centered on previously identified coordinates for tjS1 (3.8 mm lateral, 0.5 mm rostral from bregma)^24^. In each craniotomy, we made 26-32 injections of AAV-Syn-FLEX-rc[ChrimsonR-tdTomato] (Addgene 62723-AAV5; 50 nL per site; depth 250 - 300 μm; ∼1 nLs^-1^), so as to roughly cover the entire brain surface area exposed by the craniotomy. Each craniotomy was then implanted with glass cranial windows, which were made of two pieces of microscope cover glass glued together^43^. In a subsequent surgery (after >48 hours), we made small craniotomies bilaterally over ALM (1.5 lateral, 2.5 rostral from bregma) and injected rAAV2-retro-Syn-Cre (Janelia Viral Tools) at four separate depths to cover multiple cortical layers (13 nL each at 300, 500, 700, 900 μm, 52 nL total; ∼0.1 nLs^-1^). This procedure typically produced tdTomato expression under the cranial windows ∼10 days after the second surgery.

#### Intrinsic signal imaging to identify tjS1

As there can be some mouse-to-mouse variability in the exact cortical location of tjS1, we turned to intrinsic signal imaging (ISI) to functionally localize sub-regions of S1 that responded to tongue contact^24^. We expected that tjS1 thus identified was likely to be task-relevant since successful execution of our task required mice to use tactile information from the tongue. Mice were lightly anesthetized with isoflurane (0.5%–1%) and chlorprothixene (0.02 mL of 0.36 mg mL^-1^, intramuscular). With the lower jaw held open, a glass pipette attached to a piezo actuator (Q220-A4-203YB, Piezo Systems) was placed in light contact with the tip of the tongue and deflected with a half-sinusoidal waveform. ISI was performed as previously described^44,45^ using Ephus^46^. Prior to the procedure, fur around the mouth was shaved to avoid signals from inadvertent hair stimulation. In a subset of mice, we further confirmed that tongue touches were the source of the observed signals by checking that they did not appear when the piezo actuator was moved away from the tongue by ∼1 mm during stimulus delivery.

#### Craniotomies for recording

One day prior to planned silicon probe recording sessions, we made rectangular (1.5 × 2 mm) craniotomies over the target location that could accommodate multiple probe insertions. To facilitate ease of insertion, the dura was removed with sharp forceps. For the purpose of grounding the probes, a metallic screw was implanted in a smaller craniotomy over the left barrel cortex.

### Context-dependent sequence licking task

#### Task control

Mice performed sequential directed licking towards a moving lick port, as previously described^23^. A stainless steel lick port was mounted on two-axis linear motors (LSM050B-T4 and LSM025B-T4, Zaber Technologies). The height of the lick port was adjusted via a manual linear stage (MT1/M, Thorlabs). The task was controlled by a Teensy 3.1 microcontroller and custom MATLAB (R2020b) and Arduino (1.8.13) software. Lick port contacts were detected by a custom lick detection circuit (Janelia Research Campus). During sequence execution, contacts triggered port movements between locations in either a straight line or an arc. At the end of each trial, a water reward was dispensed by a water valve (LHDA0531415H, The Lee Company). The block-type at any given time was signaled to the mouse via a 470 nm LED (M470F3, Thorlabs) connected to an optic fiber (M77L01, Thorlabs) placed in front of the mouse’s face.

#### Behavioral training

Mice were first trained to execute standard licking sequences. Following an auditory cue (15 KHz pure tone), head-fixed mice made a series of five licks towards a lick port and received a water reward at the end of the sequence. We began by moving the port by small amounts from left to right after each touch. As training progressed and mice gained proficiency at modulating their lick angle (θ), we gradually increased the distance between the successive lick port positions until the first and final positions differed by 90 degrees about the midline. The lick port returned to the left side in between trials. We considered mice to be proficient at standard sequences when they could successfully target licks toward each port position with no missed licks on most trials. Once standard sequence proficiency was reached, we introduced backtracking sequences. Mice typically learned to execute backtracking sequences with <2 missed licks within 2-3 training sessions. Finally, we introduced both block-types in the same session. To enable mice to form an association between the block-type cue and backtracking expectation, the cue LED was present from the start of training. Behavioral and/or neural recordings were started after the mouse had encountered at least 10 sessions with both block-types.

#### Histology

After completion of all neural recording experiments, we perfused mice transcardially with 4% paraformaldehyde in phosphate buffered saline, and extracted the brains. Following overnight post-fixation, we obtained 100 μm coronal slices (Pelco easiSlicer, Ted Pella) and captured epifluorescence images (BX-41, Olympus). Slice images were registered to the Allen CCF and probe tracks reconstructed using SHARP-Track (github.com/cortex-lab/allenCCF)^47^.

### Quantification of behavioral variables

#### High speed videography

We captured high speed video of the mouse performing the task at 400 frames per second. Individual frames were hardware-triggered and captured bottom and side views of the mouse’s face, which was illuminated with an 850 nm LED (LED850-66-60, Roithner LaserTechnik).

Videos were acquired through a telecentric lens (0.25X, Edmund Optics), using a PhotonFocus DR1-D1312-200-G2-8 camera and Streampix 7 software (Norpix). Videos for each trial were captured separately, up to a maximum length of 20 seconds. A small minority of trials whose length exceeded this were discarded from analyses. We used DeepLabCut^48^ to fine-tune ImageNet-trained ResNet-50 convolutional neural networks (CNN) to annotate the tongue base and tongue tip in each frame.

#### Tongue kinematics

CNNs outputted low confidence for frames in which the tracked landmarks were not visible due to the tongue being inside the mouth. We considered all frames where CNN confidence exceeded 80% as frames captured during licking. For all such frames, tongue angle (θ) was measured as the angle between the vertical and the line connecting the tip and base of the tongue. By convention, negative values denoted angles to the left of the vertical (left-ward licking), while positive values denoted angles to the right. Tongue length (L) was defined as the distance between the base and the tip. To smooth out frame-to-frame jitter in the CNN outputs, we applied median and Savitzky-Golay filters across bins.

#### Lick alignment with adjusted time

The licking rates of mice varied from trial to trial. Before averaging time series of θ or L across trials or sessions, we temporally aligned individual lick events across trials as follows. First, we merged licks that occurred within 12 ms of each other and discarded licks that were shorter than 7 ms. Next, we measured the median numbers of bins comprising licks (*l*) and gaps between licks (*g*) for the entire pool of trials. Then we extracted the bins corresponding to individual licks by sampling *l/2* bins (or *(l + 1)/2* bins if *l* was odd) around the middle bin of each lick event.

Finally, we generated the adjusted time series for each trial by stitching together extracted bins for each successive lick interspersed with *g* bins of NaN (“not-a-number”) values. Trials were aligned to the middle bin of either the middle lick (Fig. 2d, Extended Data Fig. 4a-c) or the final lick of the sequence (Extended Data Fig. 4e).

In some opto trials, mice stopped licking during LED stimulation and skipped 1-2 lick cycles (Extended Data Fig. 4d). We considered a lick cycle to have been skipped if the gap between consecutive licks in the stimulation period exceeded the 99th percentile value for gap durations (*P_99_(g)*) in non-opto trials. We considered two lick cycles to have been skipped if the gap exceeded *l + 2*P_99_(g)*. Skipped lick cycles were filled in with NaN values in stitched trial data.

### Electrophysiology and spike sorting

#### Neural recordings

We used four-shank 384-channel Neuropixels 2.0 silicon probes^49^ to extracellularly record cortical neurons while mice performed the task. Prior to probe insertion into the cortex, the shanks were coated with lipophilic dye (Vybrant CM-DiI, ThermoFisher). Data were acquired at ∼30 KHz using a PXIe chassis (PXIe-1071, National Instruments) equipped with control and acquisition modules (IMEC Neuropixels PXIe control system and PXIe-8381, PXIe-6341, National Instruments), and SpikeGLX software (billkarsh.github.io/SpikeGLX).

#### Spike sorting and pre-processing

Voltage traces were processed using the Ecephys spike sorting pipeline^21^ (github.com/jenniferColonell/ecephys_spike_sorting) with Kilosort 2.5 or 3^50^. We disregarded clusters that contained spikes from multiple units or had missing spikes. This was done by discarding clusters that failed to meet any of the following thresholds for quality metrics:

I. Inter-spike interval violations (a measure of contamination by other units) < 0.5,
II. Amplitude cutoff (a measure of missing spikes) < 0.1,
III. Presence ratio (a measure of completeness of the data) > 0.8, and
IV. Contamination rate < 15%.

In a subset of sessions, cluster quality was also manually assessed via inspection in Phy (github.com/cortex-lab/phy). Across sessions, about 40% of “good” clusters identified by Kilosort passed all the cutoffs and were included in the pool of units for further analyses. For each unit, we calculated firing rates in 25 ms bins, over which we applied a Gaussian smoothing kernel in time.

#### General analysis

We used MATLAB (R2019a, Mathworks) for data pre-processing, and Python (3.11) and associated libraries for analyses, except where otherwise noted. We did not pre-determine any sample sizes. In some analyses involving trials pooled across sessions and mice, we used hierarchical bootstrap resampling as follows: to generate bootstrap samples, we randomly sampled (with replacement at each level) mice, then sessions from the sampled mice, and finally trials from the sampled sessions. We computed the relevant test statistic for each bootstrap sample. Reported confidence intervals were obtained from percentile values of these resampled test statistics.

#### Linear trial-type classifiers

To quantify differences in behavior and neural activity in different trial-types, we trained support vector machines with linear kernels (linear-SVMs) to distinguish between pairs of trial-types. We used hierarchical bootstrap resampling to generate pools of sessions; then for each iteration, we randomly sampled from the same session equal numbers of trials of each type being considered. Trials were not pooled across sessions. We report classification accuracy as the mean fraction of correctly classified trials with 10-fold cross-validation. For neural data classification, we fit linear-SVMs to distinguish between state space vectors, independently for each bin. For classification using θ, we fit linear-SVMs independently for each lick as follows. Bins corresponding to individual licks were extracted as described above. Each data-point supplied to the classifiers was then a vector of time bins of θ data for one lick.

### Optogenetic stimulation

#### LED stimulation

Optogenetic stimuli were applied using a 595 nm LED (M595F2, Thorlabs) via an optic fiber (M98L01, Thorlabs) positioned over the area of overlap of ISI signal and tdTomato expression under cranial windows. We used Wavesurfer (wavesurfer.janelia.org) to deliver 0.4 s pulses consisting of 40 Hz sinusoidal waveforms, with a 0.1 s ramp down period at the end. Because we were interested in observing trial-to-trial variability and block-type-dependent differences, we sought to avoid any ceiling effects of the optogenetic perturbations by providing the minimum light intensity required to evoke a response. In a pilot session, we started with a high LED intensity (8 mW at end of fiber) and gradually turned it down until we could no longer reliably evoke any observed effects of the optogenetic stimulation. In the following sessions where we recorded the behavior/neural data, we used ∼1 mW over this measured intensity. Additionally, we supplied randomly flashing masking light of similar wavelength, which was introduced several sessions in advance of any optogenetic interventions to enable acclimation.

#### Opto-modulation index

We measured the impact of optogenetic stimulation on firing rates of individual units as follows. Given the multisets of firing rates of a unit *u* during LED stimulation bins in non-opto (*r_u_ ^n^*) and opto (*r °*) trials, we first quantified the opto-modulation index for each unit as:

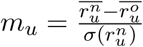

where *σ* denotes the standard deviation function. In Extended Data Fig. 4g, we plot absolute values of *m_u_*. In Extended Data Fig. 6a, we applied a symmetric log function to *m_u_*:

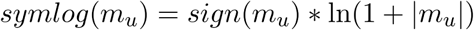

#### Quantification of perturbation magnitude

We quantified the magnitude of optogenetic perturbation of population neural activity in single opto trials as the norm of the bin-by-bin vector differences from the median in state space trajectories. We organized firing rate data in each trial *t*, consisting of *T* time bins and *N* units, into a *T* × *N* matrix *X_t_*, so that the row vectors (Τ_1_, Τ_2_, …, Τ_T_ ∈ R^N^) represented population activity vectors in each time bin. We then computed the median non-opto activity matrix *M* by considering the element-wise medians across non-opto trials. The opto-induced deviation in population activity in each opto trial was then given by the L2 norms of the row vectors of the difference matrix *M - X_t_*:

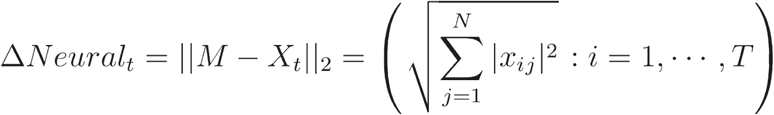

where *x_ij_* represents the element in the ith row and jth column of *X_t_*. This operation was carried out separately for opto and non-opto trials of each behavioral trial-type (Standard block standard, Backtracking block standard and Backtracking block backtracking). The resultant trial-by-trial ΔNeural time-series were then baseline-subtracted using the mean of the 200 ms of data immediately preceding onset of LED stimulation.

ΔNeural in θ-encoding subspaces (described below) was calculated similarly, but with the columns of *X_t_* containing the projections of population activity onto each θ-encoding dimension and *N* equalling the number of θ-encoding dimensions being considered (typically 4).

Δθ for each non-opto trial (Fig. 3e) was calculated simply as the difference in θ from the mean non-opto trial, averaged over bins during the second half of LED stimulation (i.e., 0.2 - 0.4 s after middle-lick). Positive values for Δθ indicated opto-induced deviations towards the left.

### θ-encoding subspaces

#### Linear decoding of θ

For each session, we identified latent dimensions, which we termed θ-encoding (θ_enc_) dimensions, that were best for linearly decoding θ. Given a *T* × *N* (*T* time bins, *N* units) neural data matrix *X*, we used Ridge regression to find a latent dimension θ_enc_ that minimized:

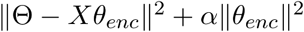

where Φ is a vector of θ values corresponding to each time bin represented by the rows of *X*, and *α* controls the strength of regularization. To generate Φ, we downsampled θ data (400 Hz) to match the time bins in *X* (40 Hz) by averaging every 10 consecutive bins. *α* was chosen on a session-by-session basis from among seven candidate values {10^e^ | e ∈ Z, −4 ≤ e ≤ 2} as the value that yielded the best 10-fold cross-validated decoding performance, measured as the coefficient of determination of predicted θ. In all places, we report decoding performance for a completely held-out set of bins (excluded from the cross-validation set used to arrive at *α*).

#### Serial deflations algorithm

We extracted a mutually orthogonal set of latent θ^(r)^_enc_ dimensions by carrying out serial deflations of each data matrix *X* (Extended Data Fig 2a). The method is an adaptation of the “Two-Block Mode A Partial Least Squares” (PLS-W2A) algorithm expounded in ref. ^51^. We began by choosing the number of rounds of deflation *R* that we wished to carry out, under the condition that *R ≤ rank(X)*. Typically this was set to 4. Then, we set *X^(0)^ ← x*. In this notation, the superscript denotes the deflation round the data correspond to. Thus *X^(0)^* is simply the original data matrix containing unit firing rates (referred to in the main text as “Full data”). Then, for each iteration *r ∈ (1, …, R)*, we performed the following steps:

1. Identify the best θ-encoding latent dimension θ^(r)^_enc_ within *X^r−1^* using Ridge regression (as outlined above).
2. Project *X^r−1^* onto θ^(r)^_enc_ to obtain scores θ^_r_ = *X^r−1^θ^(r)^_enc_*
3. Regress *X^r−1^* on θ^_r_ to obtain a 1-rank approximation of *X^r−1^*. Using ordinary least squares, we have:
4. Deflate the data by subtracting the 1-rank approximation: *X^r^* = *X^r−1^* − *X^^r−1^* The residual matrix *X^r^* is then orthogonal to θ^(r)^_enc_.

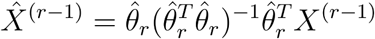

Calculation:

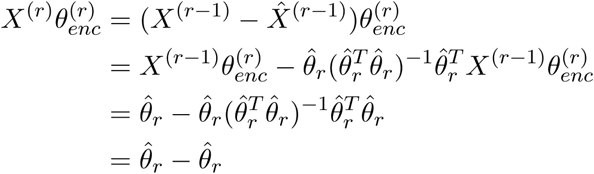
5. Increment *r* by 1. Then if *r* ≤ *R*, go to step 1, else exit.

Because of the above property of orthogonality, all θ_enc_ vectors are also mutually orthogonal. We refer to the subspace spanned by this ordered set of θ_enc_ dimensions *θ^(r)^_enc_: r = 1, …, R)* as the θ-encoding, or θ_enc_ subspace. The neural activity in the θ_enc_ space was then computed by multiplying the full data matrix *X^(0)^* with the *N* × *R* matrix whose columns are the θ_enc_ vectors, resulting in a *T* × *R* matrix. The θ_enc_ space activity of single units (shown in Fig. 1f, middle and right) was simply the sum of the approximation matrices:

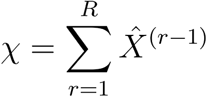

so that *X^(0)^ = x + X^(B)^* and *x^T^X^(R)^ = 0*.

#### Principal components analysis (PCA)

We used PCA either to compare principal component (PC) axes with θ_enc_ dimensions (Fig. 3b, Extended Data Fig. 2) or to visualize state space trajectories in three or two dimensions (Figs. 3f, 4a). For comparison with θ_enc_ dimensions, we used all neural data bins in the session to identify PCs. In some analyses, we computed the loss of θ decoding accuracy resulting from the removal of activity along the top PC (Extended Data Fig. 2c). To achieve this, we followed the same serial deflation algorithm outlined above, except in each round we replaced Step 1 with finding the top PC within *X^r−1^*, which then took the place of θ^(r)^_enc_. The rest of the algorithm was unaltered.

For canonical correlation analysis, we first projected data for each session onto the θ_enc_ and PC spaces (top six dimensions). Then we identified latent axes that maximized the correlation between the two activity subspaces and projected both onto those. We report the absolute Pearson correlation coefficient value between data from respective pairs of projections.

When using PCA for visualization, we identified PCs using an artificial dataset constructed by randomly choosing the same number of bins from each of the three trial-types. All trials and latent axes were then projected onto the PCs thus identified.

#### Sequence and block axes in population activity

We used linear-SVMs to identify latent axes that best discriminated pairs of trial-types. For identifying the sequence axis in each session, we trained linear-SVMs to use θ_enc_ space activity to discriminate between Backtracking block standard and Backtracking block backtracking trials, using the means of population activity in the period 0.2 to 0.5 s after the middle-lick. For identifying the block axis, we discriminated (using full data) between Standard block standard trials on the one hand, and Backtracking block standard and Backtracking block backtracking trials on the other, using the means of the 0.25 s of population activity immediately preceding the middle-lick. In both cases, trial-by-trial scores on the latent axes were obtained using leave-1-out cross-validation as follows. For each trial, we generated a unique training dataset consisting of all relevant trials in the session except the one in question. Then we fit linear-SVMs on the training data to find the latent axes, and projected data from the trial onto those axes to obtain the score. For consistency across sessions and clarity of presentation, we subtracted the corresponding intercept terms from individual trial scores to set the decision boundaries to zero. To reduce overfitting, all linear-SVM models were regularized, with the regularization parameter *C* chosen from among seven candidate values (same as for θ decoding above) as the one that yielded the best 10-fold cross-validated classification performance (the strength of regularization was inversely proportional to *C*). In cases where the trial-types being discriminated were unequally represented in the training data, *C* was multiplied by a weight that was proportionally larger for the less numerous trial-type.

#### Trial-type preference of single units

We quantified the preference of individual units for different trial-types using ideal-observer analysis. Preference for Standard versus Backtracking blocks (“Block preference”) was quantified using firing rates during the 0.25 s period prior to the middle-lick. Preference for standard versus backtracking sequences (“Sequence preference”) was quantified using firing rates during the 0.4 s period after the middle-lick. For the latter analysis, standard sequence trials from both blocks were pooled and firing rates were baseline subtracted using 0.1 s of data before the middle-lick.

In both cases, receiver operating characteristic (ROC) curves were constructed on a bin-by-bin basis with bootstrap resampling as follows. The two trial-types being compared were assigned positive or negative labels. False positive and true positive rates were then obtained by comparing those labels to positive and negative labels generated by systematically varying a criterion firing rate within data of each time bin. The area under the ROC curve (AUROC) constructed using those values represented the trial-type preference in that bin. This was repeated for each bootstrap sample. A bin was considered as showing significant preference if the 95% confidence interval of AUROC values did not include 0.5. A unit was considered as showing significant trial-type preference if there were at least 3 consecutive bins passing the significance criterion.

We also quantified the θ_enc_ space preference of individual units for licking towards the left or towards the right (Extended Data Fig. 5c). Given the θ_enc_ activity of each unit during standard sequences, we calculated the mean difference *d* between the 0.4 s window before the middle-lick (when the mouse licked left) and the 0.4 s window after the middle-lick (when the mouse licked right). The left licking preference was then given by the symmetric log of *d*:

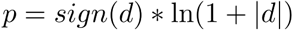

### Population dynamics

#### Neural population activity parameters

To quantify the nature of neural dynamics in each trial, we measured a number of parameters of population activity in the 0.25 s window before the middle-lick (corresponding to the same time bins as were used for the block axis calculation above), as follows. Unit σ was quantified as the standard deviation of firing rates across bins for each unit, averaged across units. Population correlation was calculated as the correlation coefficient in firing rates between each pair of units, averaged across all pairs of units. Temporal cosines were the vector cosine similarities between population activity vectors in consecutive time bins, averaged across all pairs of bins. Trial drift was quantified as the L2-norms of the bin-by-bin vector differences from the median data matrix, averaged across bins.

#### Random Forest classifiers

We trained Random Forest classifiers to classify the block-type of individual trials using block axis scores and the above activity parameters as features. For each hierarchical bootstrap sample, values for all the features were z-scored, and a random 90% of the trials were used to train classifiers. Model performance, quantified as the proportion of correctly classified trials, was then measured on the held-out test set. To reduce overfitting, we made only 60% of training trials available to individual trees and prohibited leaf nodes with fewer than 20 trials. To test their importance for classification performance, individual features were randomly shuffled. References

---

**Extended Data Fig. 1.**
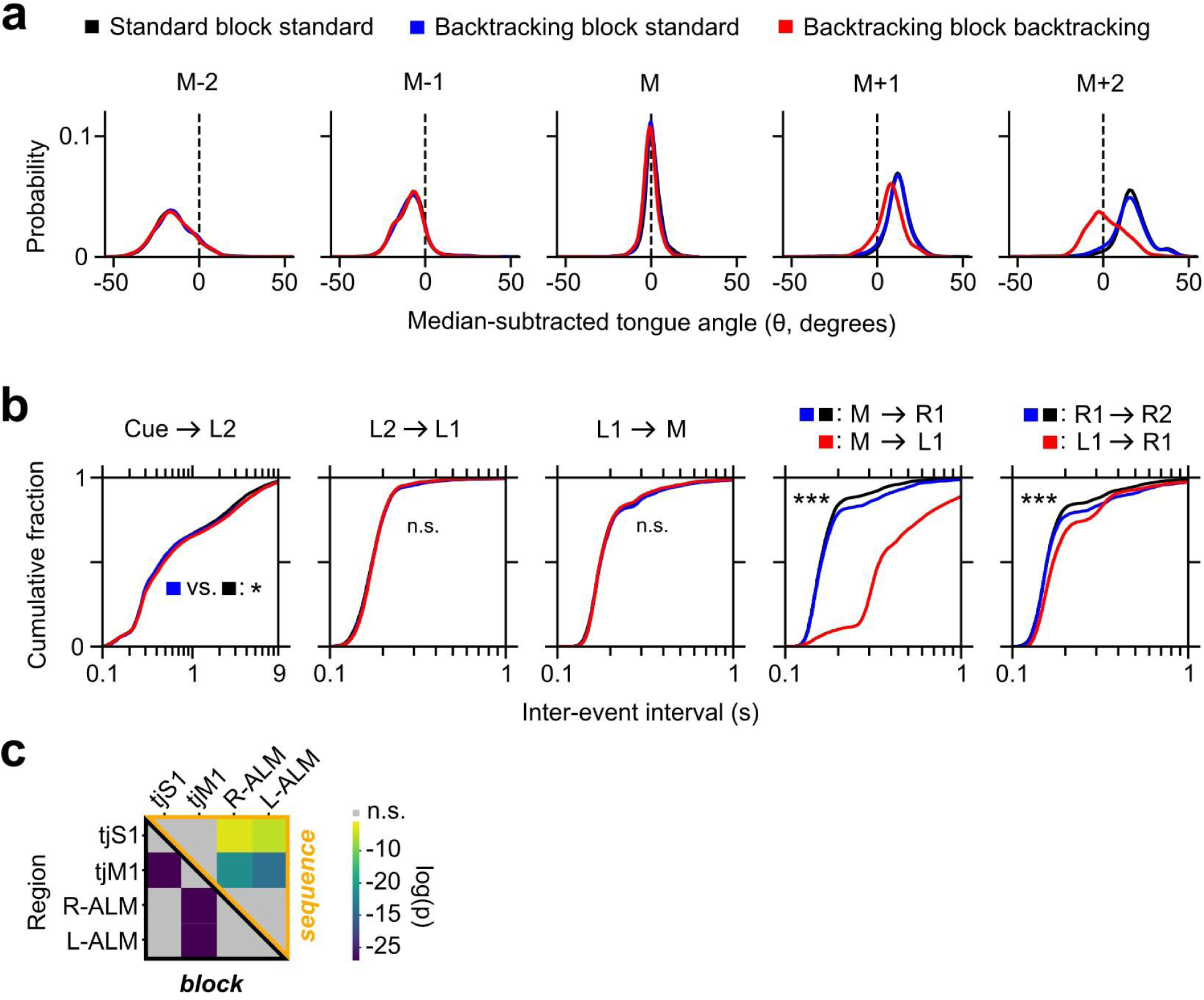
Subtle block-dependent differences in licking rate **a,** Kernel density estimate plots of lick angle for the middle-lick (M) and neighboring licks during execution of Standard block standard (black), Backtracking block standard (blue) and Backtracking block backtracking (red) trials. For each individual lick, tongue angle was considered when tongue was at maximum protrusion and the median middle-lick value for the session was subtracted. **b,** Cumulative histograms of time intervals between successive pairs of events, in different trial-types with the same color scheme as in (a). Schematics above the plots indicate the relevant pairs of events for each plot and trial-type. Two-sided Kolmogorov-Smirnov test, *** p < 0.001, * p < 0.05. **c,** Heatmap of p-values testing the null hypothesis that the median trial-type classification performance values from Figure 1g are the same between each pair of regions. Wilcoxon signed rank test with Bonferroni correction for multiple comparisons.

**Extended Data Fig. 2.**
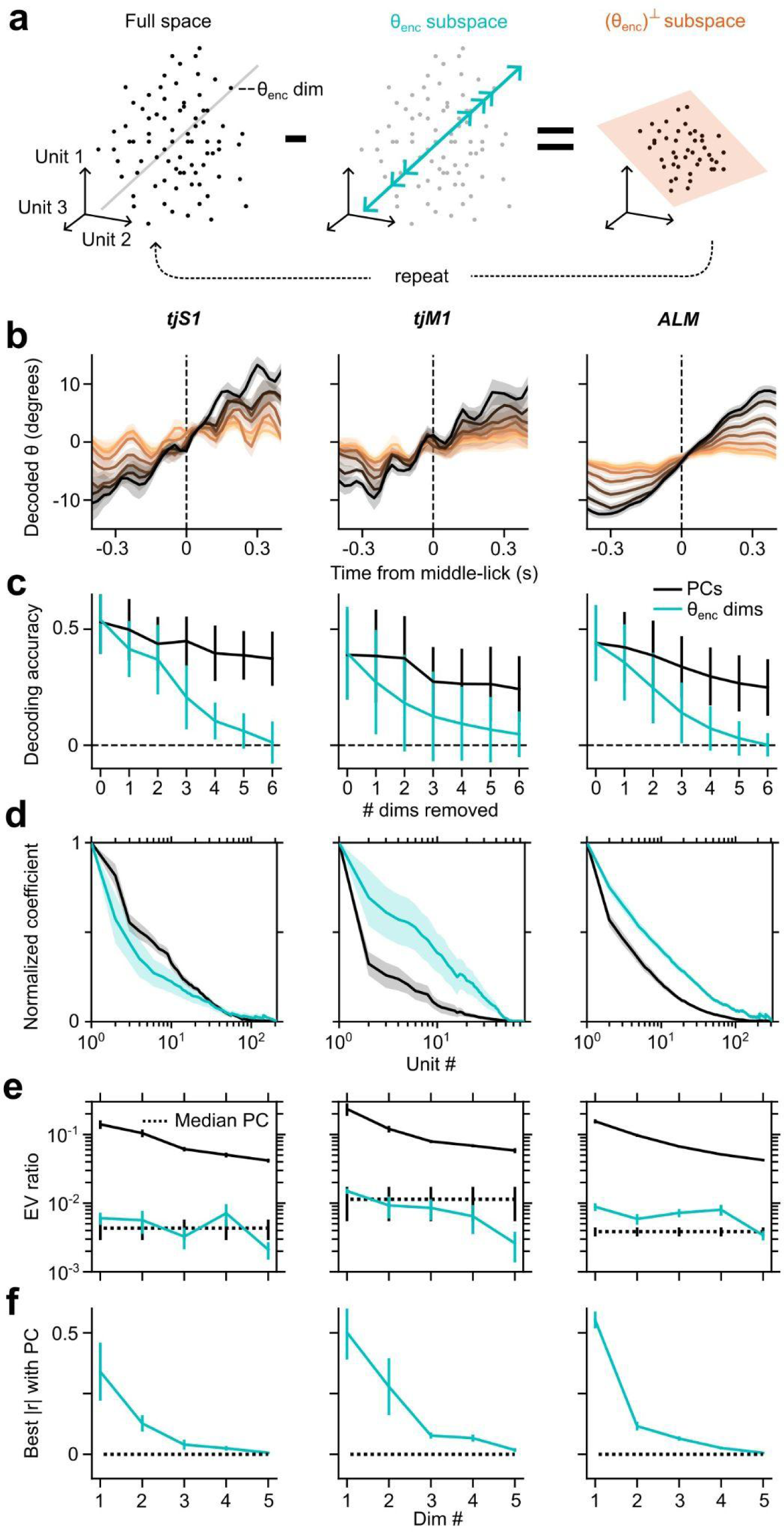
Exploration of θ_enc_ dimensions. **a,** Schematic of process for identifying and extracting an orthogonal set of θ_enc_ dimensions by serial deflations of the data. **b,** Linearly decoded θ from neural activity in different cortical regions after successive rounds of θ_enc_ deflations Colors from black to tan indicate the number of θ_enc_ dimensions removed from the data, with black being zero. **c,** θ decoding accuracy versus number of principal components (black) or θ_enc_ dimensions (cyan) removed. Data are mean ± s.d. **d,** Normalized loadings of individual units for the first PC (black) or θ_enc_ dimension (cyan). **e,** Explained variance ratios of the top PCs, θ_enc_ dimensions and median PCs. **f,** Canonical correlation r values between the top θ_enc_ dimensions and the corresponding PCs. Dotted line shows the same comparison between median and top PCs. All data are mean ± s.e.m. unless otherwise noted.

**Extended Data Fig. 3.**
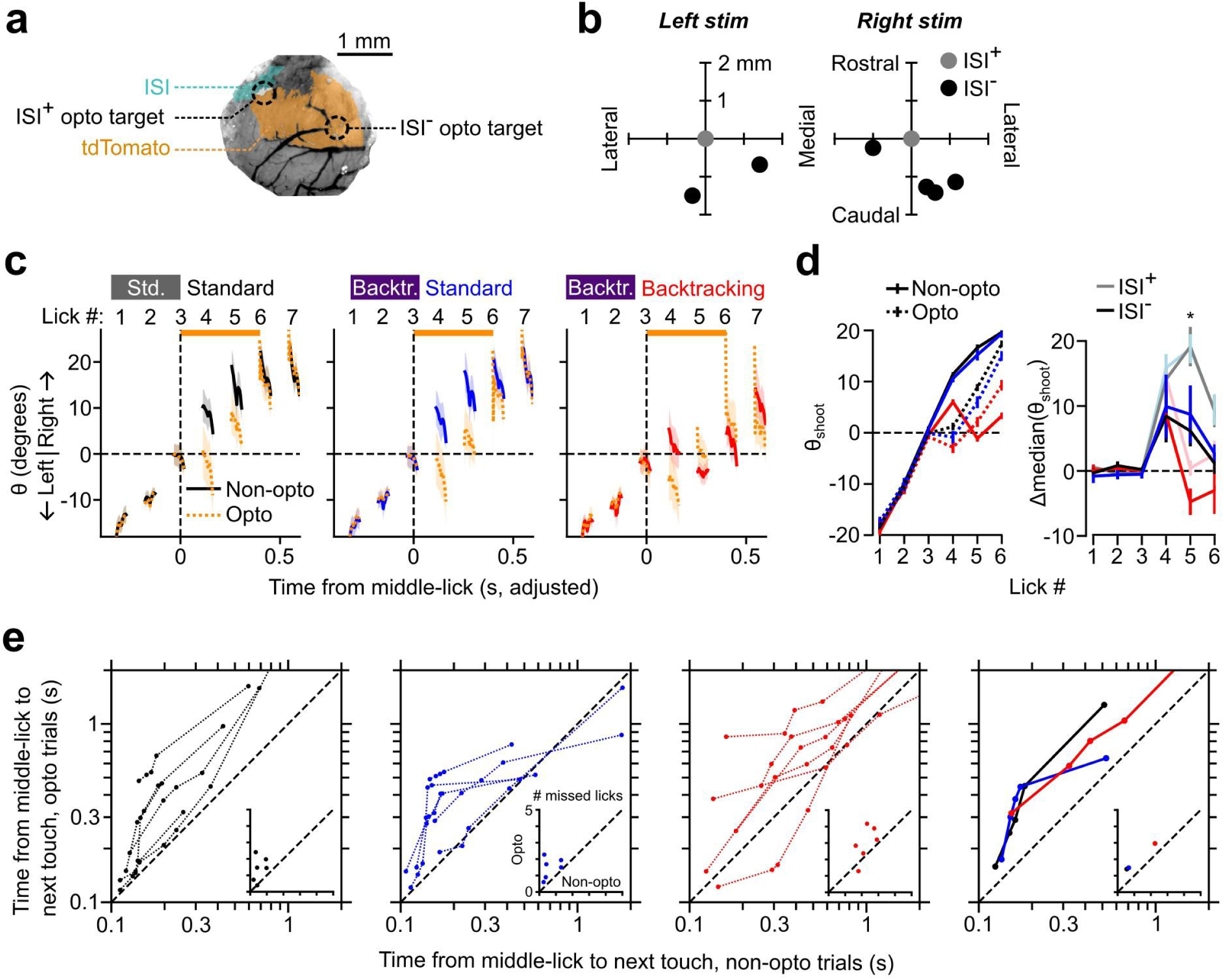
(related to Fig. 2) **a,** Example image of the same cranial window as in Fig. 2a, showing in addition the target location for optogenetic stimulation in a region lacking ISI signal (ISI**^-^**). **b,** Target locations for optogenetic stimulation in ISI**^-^** sessions relative to ISI^+^ sessions. **c,** θ versus time from middle-lick for the three trial types in sessions with ISI**^-^** opto-stimulation. Orange bar denotes time of opto-stimulation. Data are mean ± s.e.m. **d,** Left, θ_shoot_ for each lick in non-opto and opto trials, in sessions with ISI**^-^** opto-stimulation. Right, difference in median θ_shoot_ values between non-opto and opto trials. For comparison, data from ISI^+^ sessions (Fig. 2e) are reproduced. Data are mean ± s.e.m. **e,** Quantile-quantile plots showing time between middle-lick and next touch for opto versus non-opto trials, in sessions with ISI**^-^** opto-stimulation. The left three plots show values for the three trial-types in individual sessions, the right plot shows medians across sessions. Insets are scatter-plots showing the mean number of missed licks between middle-lick and next touch for opto versus non-opto trials. Wilcoxon signed rank test, * p < 0.5.

**Extended Data Fig. 4.**
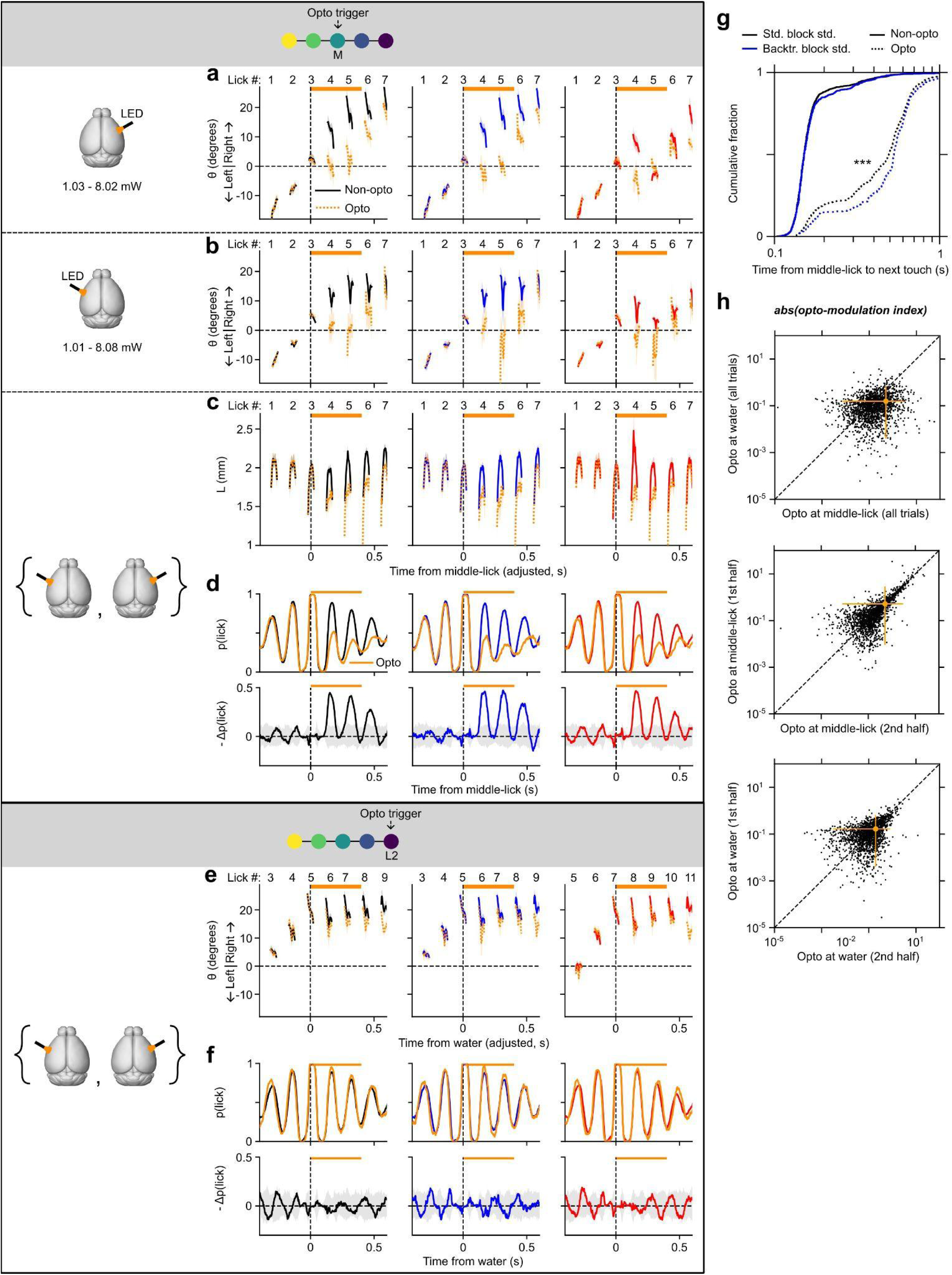
(related to Fig. 2) **a-d,** Opto-stimulation triggered by touch at middle position. **a,** θ versus time from middle-lick for sessions where the stimulus LED was positioned over right tjS1. Solid lines represent non-opto trials: black lines represent Standard block standard trials, blue represents Backtracking block standard trials and red represents Backtracking block backtracking trials. Orange dotted lines represent the corresponding opto trials. Orange bar denotes time of opto-stimulation. The same color scheme is followed in the other sub-figures. The range of light intensities at the optic fiber tip are indicated. Data are mean ± s.e.m. **b,** Same as (a) but for sessions where opto-LED was positioned over left tjS1. **c,** Tongue length versus time from middle-lick for different trial types. Data are mean ± s.e.m. **d,** Top row, probability of the tongue being out of the mouth versus time from middle lick for the different trial types. Bottom row, negative change in lick probability produced by the opto-stimulation. The gray shaded areas show 95% confidence intervals for null distributions generated by shuffling opto and non-opto trials. **e-f,** Opto-stimulation triggered by touch at final position (i.e., water trigger). **e,** θ versus time from water for different trial types with the same color scheme as (a-c). **f,** Same as (d) but plotted against time from water on the x-axis. **g,** Cumulative histograms showing time between middle-lick and next touch for Standard block standard and Backtracking block standard, non-opto and opto trials. Two-sided Kolmogorov-Smirnov test, *** p < 0.001. **h,** Scatter-plots of opto-modulation indices for individual units. Top, opto-modulation indices for stimulation at water versus middle-lick. Middle, opto-modulation indices for a random half of stimulation at middle-lick trials versus the same for the other half. Bottom, same as middle but for trials with opto-stimulation at water. Orange markers and lines show means and 95% confidence intervals respectively.

**Extended Data Fig. 5.**
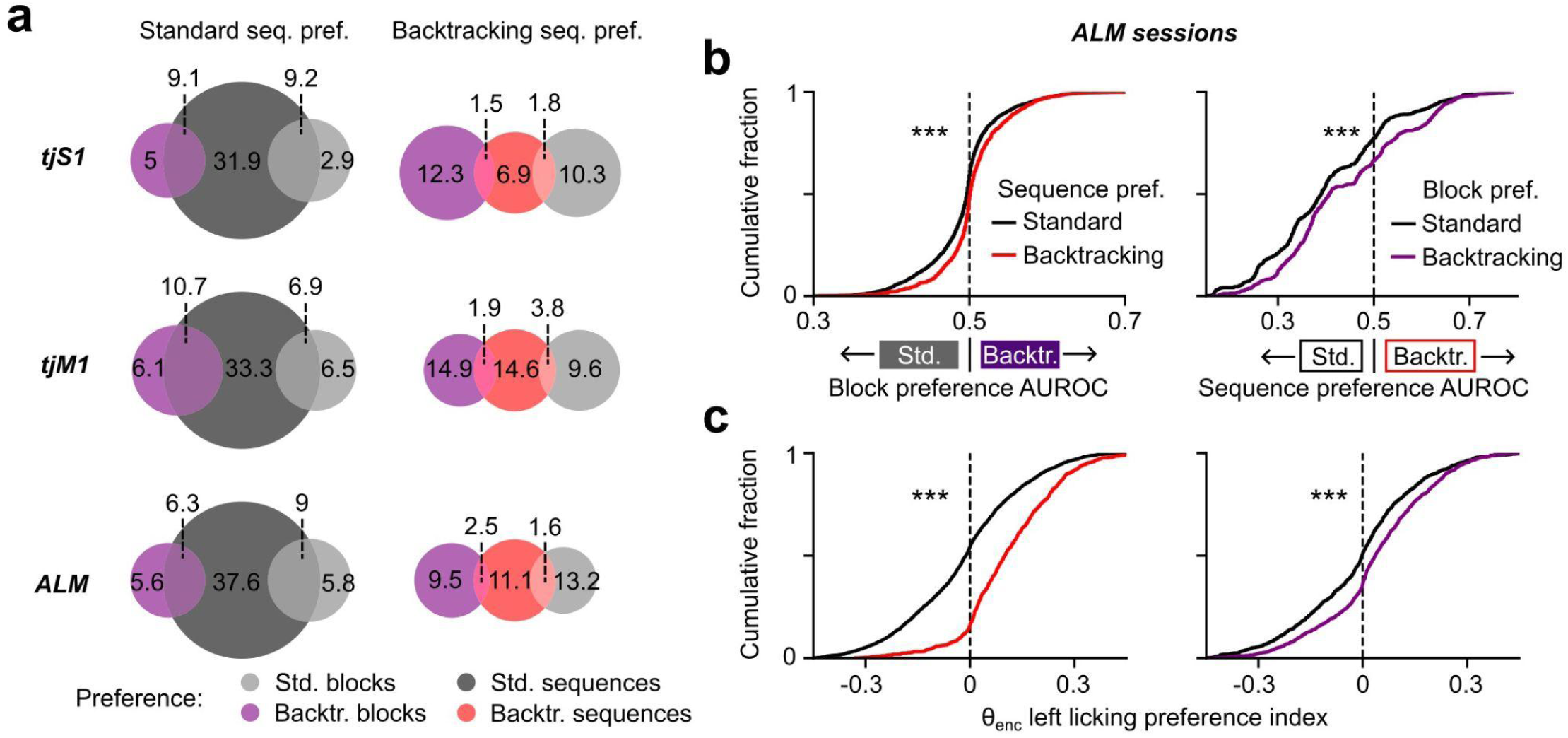
Preference of single units for different blocks and/or sequences **a,** Venn diagrams of percentages of units in each cortical region showing preferences for either block or sequence type. Left column, overlap of units preferring standard sequences with units preferring either block type. Right column, overlap of units preferring backtracking sequences with units preferring either block type. Note that units showing preference for a given block are represented twice in each row. **b,** Left, cumulative histograms of block preference AUROC values for units showing preference for either sequence type. Right, cumulative histograms of sequence preference AUROC values for units showing preference for either block type. **c,** Left, cumulative histograms of left licking preference in θ_enc_ activity for units showing preference for either sequence type. Right, same as left but for units showing block type preference. Two-sided Kolmogorov-Smirnov test, *** p < 0.001.

**Extended Data Fig. 6.**
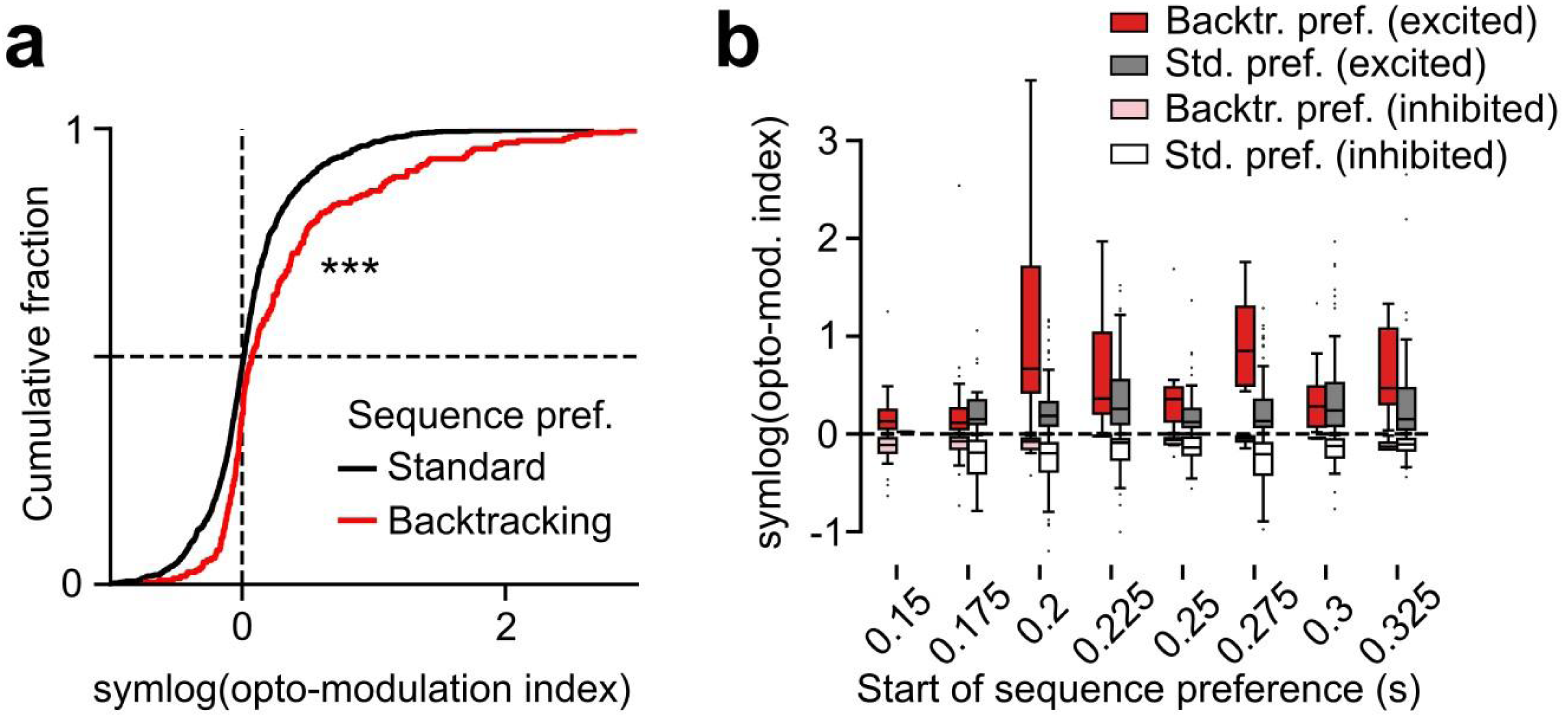
Backtracking sequence-preferring units were more excited by opto-stimulation a, Cumulative histograms showing symmetric logs of opto-modulation indices for single units (Methods) that prefer either standard or backtracking sequences. **b,** Box-plots showing symmetric logs of opto-modulation indices for individual units, divided into groups based on the time after middle-lick at which they started showing a preference for either sequence-type. For each such group, data are then shown separately for standard- or backtracking-preferring units that were excited or inhibited by the opto-stimulation. Two-sided Kolmogorov-Smirnov test, *** p < 0.001.

**Extended Data Fig. 7.**
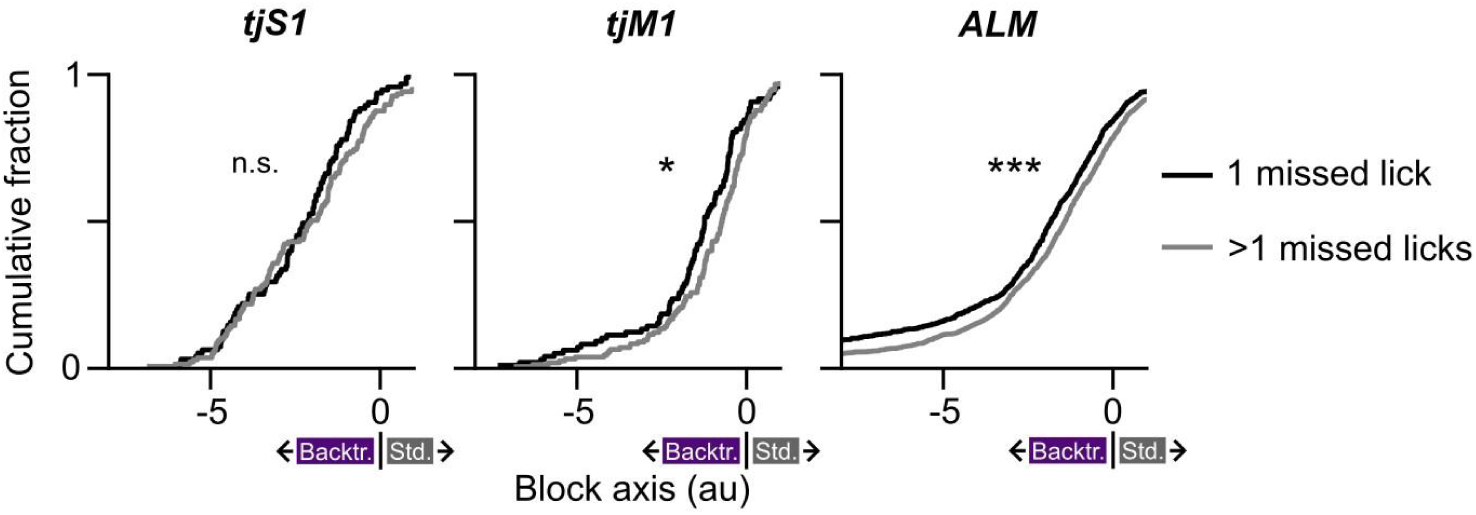
(related to Fig. 4) Cumulative histograms of block axis projections for non-opto backtracking trials in which mice required either a single or multiple missed licks to successfully execute backtracking. Two-sided Kolmogorov-Smirnov test, *** p < 0.001, * p < 0.05.

**Extended Data Fig. 8.**
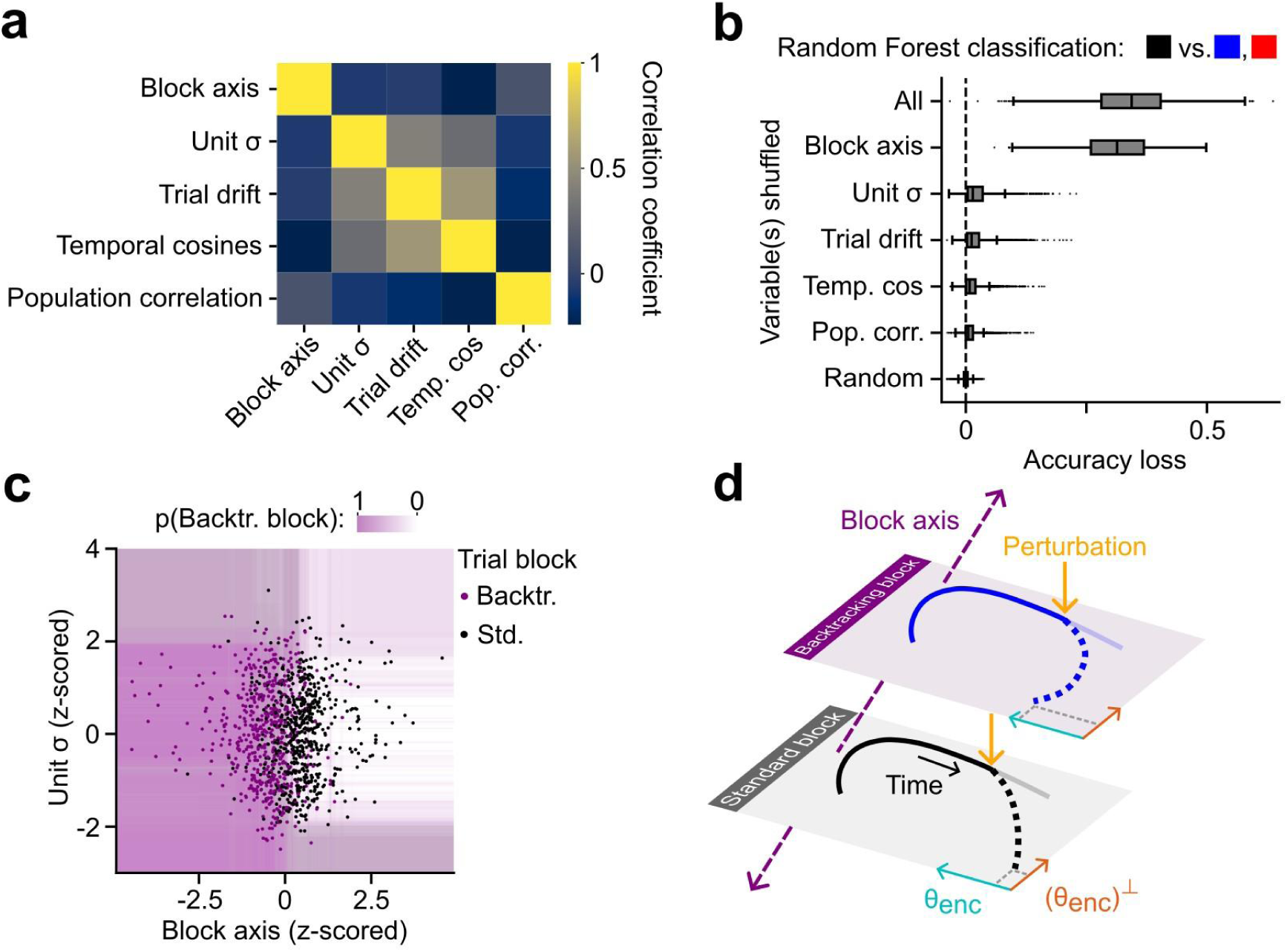
Trials from the two blocks showed similar dynamics **a,** Heatmap of Pearson correlation coefficients between the five features of neural dynamics quantified for each trial (see Methods). **b,** Loss in binary block type classification accuracy by Random Forest classifiers due to shuffling all features or single features. **c,** Scatter-plot of Unit σ versus block axis projections for a held out set of Standard and Backtracking block trials. The background heatmap shows probability of classification as a Backtracking block trial by any given tree in a Random Forests ensemble trained to classify block type. **d,** Schematic of neural dynamical model.

## References

1. Perich, M. G. et al. Motor cortical dynamics are shaped by multiple distinct subspaces during naturalistic behavior. Preprint at 10.1101/2020.07.30.228767 (2020).

2. Scott, S. H., Cluff, T., Lowrey, C. R. & Takei, T. Feedback control during voluntary motor actions. Curr. Opin. Neurobiol. 33, 85–94 (2015).

3. Tanji, J. & Shima, K. Role for supplementary motor area cells in planning several movements ahead. Nature 371, 413–416 (1994).

4. Pruszynski, J. A. et al. Primary motor cortex underlies multi-joint integration for fast feedback control. Nature 478, 387–390 (2011).

5. Scott, S. H. A Functional Taxonomy of Bottom-Up Sensory Feedback Processing for Motor Actions. Trends Neurosci. 39, 512–526 (2016).

6. Desrochers, T. M., Burk, D. C., Badre, D. & Sheinberg, D. L. The Monitoring and Control of Task Sequences in Human and Non-Human Primates. Front. Syst. Neurosci. 9, (2016).

7. Stavisky, S. D., Kao, J. C., Ryu, S. I. & Shenoy, K. V. Motor Cortical Visuomotor Feedback Activity Is Initially Isolated from Downstream Targets in Output-Null Neural State Space Dimensions. Neuron 95, 195–208.e9 (2017).

8. Sauerbrei, B. A. et al. Cortical pattern generation during dexterous movement is input-driven. Nature 577, 386–391 (2020).

9. Inagaki, H. K. et al. Neural Algorithms and Circuits for Motor Planning. Annu. Rev. Neurosci. 45, 249–271 (2022).

10. Yang, W., Tipparaju, S. L., Chen, G. & Li, N. Thalamus-driven functional populations in frontal cortex support decision-making. Nat. Neurosci. 25, 1339–1352 (2022).

11. Codol, O. et al. Sensorimotor feedback loops are selectively sensitive to reward. eLife 12, e81325 (2023).

12. Michaels, J. A. et al. Sensory expectations shape neural population dynamics in motor circuits. Preprint at 10.1101/2024.12.22.629295 (2024).

13. Pruszynski, J. A. & Scott, S. H. Optimal feedback control and the long-latency stretch response. Exp. Brain Res. 218, 341–359 (2012).

14. Reschechtko, S. & Pruszynski, J. A. Stretch reflexes. Curr. Biol. 30, R1025–R1030 (2020).

15. Gao, Z. et al. A cortico-cerebellar loop for motor planning. Nature 563, 113–116 (2018).

16. Churchland, M. M. & Shenoy, K. V. Temporal Complexity and Heterogeneity of Single-Neuron Activity in Premotor and Motor Cortex. J. Neurophysiol. 97, 4235–4257 (2007).

17. Churchland, M. M. et al. Neural population dynamics during reaching. Nature 487, 51–56 (2012).

18. Xue, C., Kramer, L. E. & Cohen, M. R. Dynamic task-belief is an integral part of decision-making. Neuron 110, 2503–2511.e3 (2022).

19. Wang, S., Falcone, R., Richmond, B. & Averbeck, B. B. Attractor dynamics reflect decision confidence in macaque prefrontal cortex. Nat. Neurosci. 26, 1970–1980 (2023).

20. Bollu, T. et al. Cortex-dependent corrections as the tongue reaches for and misses targets. Nature 594, 82–87 (2021).

21. Chen, S. et al. Brain-wide neural activity underlying memory-guided movement. Cell 187, 676–691.e16 (2024).

22. Li, N., Chen, T.-W., Guo, Z. V., Gerfen, C. R. & Svoboda, K. A motor cortex circuit for motor planning and movement. Nature 519, 51–56 (2015).

23. Xu, D. et al. Cortical processing of flexible and context-dependent sensorimotor sequences. Nature 603, 464–469 (2022).

24. Mayrhofer, J. M. et al. Distinct Contributions of Whisker Sensory Cortex and Tongue-Jaw Motor Cortex in a Goal-Directed Sensorimotor Transformation. Neuron 103, 1034–1043.e5 (2019).

25. Meirhaeghe, N., Riehle, A. & Brochier, T. Parallel movement planning is achieved via an optimal preparatory state in motor cortex. Cell Rep. 42, 112136 (2023).

26. Sadtler, P. T. et al. Neural constraints on learning. Nature 512, 423–426 (2014).

27. Chaudhuri, R., Gerçek, B., Pandey, B., Peyrache, A. & Fiete, I. The intrinsic attractor manifold and population dynamics of a canonical cognitive circuit across waking and sleep. Nat. Neurosci. 22, 1512–1520 (2019).

28. Nieh, E. H. et al. Geometry of abstract learned knowledge in the hippocampus. Nature 595, 80–84 (2021).

29. Stringer, C. et al. Spontaneous behaviors drive multidimensional, brainwide activity. Science 364, eaav7893 (2019).

30. Musall, S., Kaufman, M. T., Juavinett, A. L., Gluf, S. & Churchland, A. K. Single-trial neural dynamics are dominated by richly varied movements. Nat. Neurosci. 22, 1677–1686 (2019).

31. Shenoy, K. V., Sahani, M. & Churchland, M. M. Cortical Control of Arm Movements: A Dynamical Systems Perspective. Annu. Rev. Neurosci. 36, 337–359 (2013).

32. Ito, B. S., Gao, Y., Kardon, B. & Goldberg, J. H. A collicular map for touch-guided tongue control. Nature 637, 1143–1151 (2025).

33. Zimnik, A. J. & Churchland, M. M. Independent generation of sequence elements by motor cortex. Nat. Neurosci. 24, 412–424 (2021).

34. Eriksson, D., Heiland, M., Schneider, A. & Diester, I. Distinct dynamics of neuronal activity during concurrent motor planning and execution. Nat. Commun. 12, 5390 (2021).

35. Kaufman, M. T., Churchland, M. M., Ryu, S. I. & Shenoy, K. V. Cortical activity in the null space: permitting preparation without movement. Nat. Neurosci. 17, 440–448 (2014).

36. Churchland, M. M. & Shenoy, K. V. Preparatory activity and the expansive null-space. Nat. Rev. Neurosci. 25, 213–236 (2024).

37. Han, W. et al. Integrated Control of Predatory Hunting by the Central Nucleus of the Amygdala. Cell 168, 311–324.e18 (2017).

38. Kawai, R. et al. Motor Cortex Is Required for Learning but Not for Executing a Motor Skill. Neuron 86, 800–812 (2015).

39. Mizes, K. G. C., Lindsey, J., Escola, G. S. & Ölveczky, B. P. Dissociating the contributions of sensorimotor striatum to automatic and visually guided motor sequences. Nat. Neurosci. 26, 1791–1804 (2023).

40. Merel, J., Botvinick, M. & Wayne, G. Hierarchical motor control in mammals and machines. Nat. Commun. 10, (2019).

41. Mizes, K. G. C., Lindsey, J., Escola, G. S. & Ölveczky, B. P. The role of motor cortex in motor sequence execution depends on demands for flexibility. Nat. Neurosci. (2024) doi:10.1038/s41593-024-01792-3.

42. Heindorf, M., Arber, S. & Keller, G. B. Mouse Motor Cortex Coordinates the Behavioral Response to Unpredicted Sensory Feedback. Neuron 99, 1040–1054.e5 (2018).

43. Kwon, S. E., Yang, H., Minamisawa, G. & O’Connor, D. H. Sensory and decision-related activity propagate in a cortical feedback loop during touch perception. Nat. Neurosci. 19, 1243–1249 (2016).

44. O’Connor, D. H. et al. Neural coding during active somatosensation revealed using illusory touch. Nat. Neurosci. 16, 958–965 (2013).

45. Minamisawa, G., Kwon, S. E., Chevée, M., Brown, S. P. & O’Connor, D. H. A Non-canonical Feedback Circuit for Rapid Interactions between Somatosensory Cortices. Cell Rep. 23, 2718–2731.e6 (2018).

46. Suter, B. Ephus: multipurpose data acquisition software for neuroscience experiments. Front. Neural Circuits 4, (2010).

47. Shamash, P., Carandini, M., Harris, K. D. & Steinmetz, N. A. A tool for analyzing electrode tracks from slice histology. Preprint at 10.1101/447995 (2018).

48. Mathis, A. et al. DeepLabCut: markerless pose estimation of user-defined body parts with deep learning. Nat. Neurosci. 21, 1281–1289 (2018).

49. Steinmetz, N. A. et al. Neuropixels 2.0: A miniaturized high-density probe for stable, long-term brain recordings. Science 372, eabf4588 (2021).

50. Pachitariu, M., Sridhar, S., Pennington, J. & Stringer, C. Spike sorting with Kilosort4. Nat. Methods 21, 914–921 (2024).

51. Wegelin, J. A. A Survey of Partial Least Squares (PLS) Methods, with Emphasis on the Two-Block Case. https://stat.uw.edu/sites/default/files/files/reports/2000/tr371.pdf (2000).

